# Structures of the interleukin 11 signalling complex reveal dynamics of gp130 extracellular domains and the inhibitory mechanism of a cytokine variant

**DOI:** 10.1101/2022.07.21.500383

**Authors:** Riley D. Metcalfe, Eric Hanssen, Ka Yee Fung, Kaheina Aizel, Clara C. Kosasih, Courtney O. Zlatic, Larissa Doughty, Craig J. Morton, Andrew P. Leis, Michael W. Parker, Paul R. Gooley, Tracy L. Putoczki, Michael D.W. Griffin

## Abstract

Interleukin (IL-)11, an IL-6 family cytokine, has pivotal roles in numerous autoimmune diseases, fibrotic complications, and solid cancers. Despite intense therapeutic targeting efforts, structural understanding of IL-11 signalling and mechanistic insights into current inhibitors is lacking. Here we present cryo-EM and crystal structures of the IL-11 signalling complex, including the complex containing the complete extracellular domains of the shared IL-6 family β-receptor, gp130. We show that the membrane-proximal domains of gp130 are dynamic and do not participate in complex assembly. We demonstrate that the cytokine mutant ‘IL-11 Mutein’ competitively inhibits signalling in human cell lines. Structural shifts in IL-11 Mutein underlie inhibitory activity by altering cytokine binding interactions at all three receptor-engaging sites and abrogating the final gp130 binding step. Our results reveal the structural basis of IL-11 signalling, define the molecular mechanisms of an inhibitor, and advance understanding of gp130-containing receptor complexes, with potential applications in therapeutic development.

## Introduction

Interleukin (IL-)11 is secreted by numerous immune cells including CD8+ T-cells, B-cells, macrophages, natural killer (NK) cells, γδT cells, and eosinophils, and has been implicated in the differentiation of B-cells, T-cells, and anti-tumour immune responses^1-5^. IL-11 is also produced by inflammatory fibroblasts and epithelial cells resulting in a coordinated wound response to enable the maintenance of mucosal barriers^6-11^. We now appreciate that the biological function of IL-11 spans beyond its classical role in megakaryocytopoiesis^12,13^, where recombinant IL-11 (Neumega) is FDA approved to support platelet reconstitution following chemotherapy^14^. Critical pathological roles for dysregulated IL-11 have been identified in autoimmune diseases including arthritis, asthma, inflammatory bowel disease, multiple sclerosis, and systemic sclerosis^3,4,6,7,15,16^. In addition, IL-11 drives fibrotic complications of the gastrointestinal tract, heart, kidney, liver and lung^9-11,17-21^, and promotes the growth of several malignancies, including breast, lung, endometrial and gastrointestinal cancers^22-26^. Despite these emerging physiological and pathological roles, structural understanding of IL-11 signalling has remained limited.

IL-11 activates downstream signalling pathways by binding its two cell surface receptors; the IL-11 specific receptor, IL-11Rα, and the signal-transducing receptor, glycoprotein (gp)130^13,27^. Following the formation of the receptor complex, the Janus kinase (JAK)/signal transducer and activator of stat (STAT) and extracellular signal regulated kinase (ERK)/mitogen-activated protein kinase (MAPK) pathways are primarily activated^13,28-30^. IL-11 is a member of the IL-6 family of cytokines, which also includes IL-6, leukemia inhibitory factor (LIF), oncostatin M (OSM), ciliary neurotrophic factor (CNTF), cardiotrophin-1 (CT-1), cardiotrophin-like cytokine (CLC), IL-27, IL-31, IL-35, IL-39 and an analogue of IL-6 from human herpes virus 8 (viral IL-6, vIL-6)^31-33^. The IL-6 family is commonly defined by the shared use of gp130, and thus overlaps with the IL-12 family^34^. IL-11 and IL-6 utilise a homodimer of gp130^27,35,36^ while other IL-6 family members exploit a heterodimer of gp130 with a second signal-transducing receptor, such as LIF receptor (LIFR), OSM receptor (OSMR), IL-27R/WSX-1 or IL-12 receptor β-2 subunit (IL-12Rβ2)^37^. IL-31 is unique in its use of the gp130-like receptor chain, IL-31RA, and OSMR. gp130 is a member of the ‘tall’ type-I cytokine receptor family, with a large extracellular region comprising six domains, D1-D6^28^. Crystal structures of IL-6^38^, vIL-6^35^, and LIF in complex with D1-D3 show that these membrane distal domains are involved in cytokine binding. However, the nature of the membrane proximal D4-D6 domains of gp130 within cytokine-receptor complexes has not been fully elucidated.

Inhibition of IL-11 signalling has been shown to provide therapeutic benefit in models of arthritis, multiple sclerosis, neointimal hyperplasia, multiple fibrotic diseases, and gastrointestinal cancers^9-11,13,17,21,22,25,39,40^. Current IL-11 signalling inhibitors include IL-11 mutants^41,42^ and antibodies against either IL-11^9,21,43^ or IL-11Rα^11,44-46^. However, mechanistic understanding of their modes of action is limited in the absence of detailed molecular understanding of the IL-11 signalling complex. Informed development of new and existing signalling inhibitors requires a comprehensive understanding of the structure and assembly mechanisms of the IL-11 signalling complex.

Here, we present structures of the IL-11 signalling complex, providing exquisite detail of the molecular mechanisms of complex formation and the structure and dynamics of the complete extracellular domains of gp130 within the complex. We characterise an IL-11 variant, “IL-11 Mutein”, that potently inhibits IL-11 signalling and describe the detailed mechanism of its action. Our results validate IL-11 Mutein as a tool IL-11 signalling inhibitor and provide a facile method for its production. Our insights reveal the structural basis of IL-11 signalling and provide invaluable molecular platforms for development of existing and novel therapeutics targeting IL-11 signalling and other class I cytokines.

## Results

### The structure of the IL-11 signalling complex

To understand the molecular mechanisms underpinning IL-11 signalling complex formation, we solved three structures of the complex (Figure 1) containing either the cytokine binding domains of gp130 (gp130_D1-D3_) or the complete extracellular domains of gp130 (gp130_EC_) using electron cryo-microscopy (cryo-EM) and X-ray crystallography. Complexes comprised an N-terminally truncated form of IL-11 (IL-11_Δ10_) or full length IL-11 (IL-11_FL_), and a C-terminally truncated form of IL-11Rα (IL-11Rα_D1-D3_) or the complete extracellular domains of IL-11Rα (IL-11Rα_EC_) described previously^47^.

**Figure 1:**
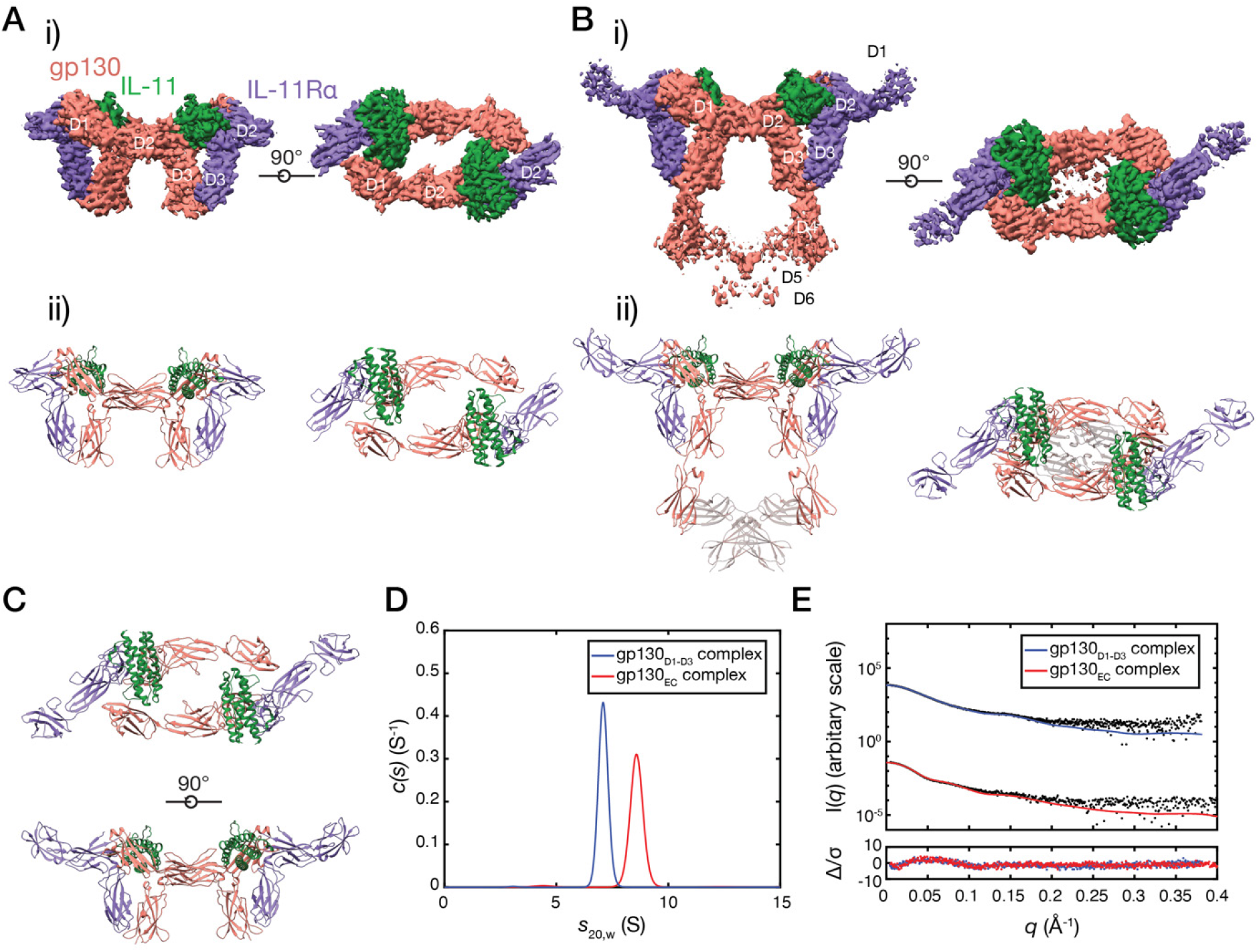
Structure of the IL-11 signalling complex (IL-11, green; IL-11Rα, purple; gp130 salmon). A) Cryo-EM density map, i), and atomic model, ii) of the IL-11_Δ10_/IL-11Rα_D1-D3_/gp130_D1-D3_ complex. D1 of IL-11Rα was not modeled in this structure. B) Cryo-EM density map i) and atomic model, ii) of the IL-11_Δ10_/IL-11Rα_D1-D3_/gp130_EC_ complex. The position of the D5-D6 domains of gp130 is indicated as transparent ribbons. These domains were not included in the deposited model. Individual domains of IL-11Rα and gp130 are indicated on the cryoEM maps. C) Crystal structure of the IL-11_FL_/IL-11Rα_EC_/gp130_D1-D3_ complex showing one hexamer of the asymmetric unit. D) Continuous sedimentation coefficient (c(s)) distributions for the complexes formed between i) IL-11_Δ10_/IL-11Rα_D1-D3_/gp130_D1-D3_ and IL-11_Δ10_/IL-11Rα_D1-D3_/gp130_EC_. E) SAXS data for the IL-11_Δ10_/IL-11Rα_D1-D3_/gp130_D1-D3_ complex and the IL-11_Δ10_/IL-11Rα_D1-D3_/gp130_EC_ complex.

We obtained a 3.5 Å resolution cryo-EM reconstruction of the IL-11_Δ10_/IL-11Rα_D1-D3_/gp130_D1-D3_ complex (referred to as the gp130_D1-D3_ complex) (Figure 1Ai), and a 3.76 Å resolution cryo-EM reconstruction of the IL-11_Δ10_/IL-11Rα_D1-D3_/gp130_EC_ complex (referred to as the gp130_EC_ complex) (Figure 1Bi, Supplementary Figure 1A-D, Supplementary Table 1). We used these data to build and refine the atomic models of the IL-11 signalling complex (Figure 1Aii-Bii). We also obtained crystals of the IL-11_FL_/IL-11Rα_EC_/gp130_D1-D3_ complex that diffracted anisotropically to 3.8 Å. We used the cryoEM structure of the gp130_D1-D3_ complex to phase the X-ray diffraction data by molecular replacement (Figure 1C). Three IL-11 signalling complexes were arranged in a triangular configuration in the asymmetric unit providing 6-fold non-crystallographic symmetry to aid refinement (Supplementary Figure 2A-C).

Secondary structure was clearly visible in both cryoEM density maps, larger side chains were generally defined, the α-helical structure of the cytokine was clear, and β-strands were generally separated, consistent with maps reconstructed at these resolutions (Supplementary Figure 1B, D). D1 of IL-11Rα was not visible in the gp130_D1-D3_ complex map and was poorly defined in the gp130_EC_ complex map (Supplementary Figure 1Di), suggesting this domain is flexible. Satisfactory density defining the positions of the β-sheets of IL-11Rα D1 facilitated refinement of the domain in the gp130_EC_ complex and the crystal structure, confirming the average positions of this domain in the complex (Supplementary Figure 1B,D, Supplementary Figure 2). N-linked glycans were visible on IL-11Rα (N105, N172) and gp130 (N21, N61, N135) in all structures. D5-D6 of gp130 were poorly defined in the gp130_EC_ complex density (Figure 1Bi) and were not included in the deposited model. However, density was sufficient to refine the positions of these domains as rigid-bodies (shown in grey in Figure Bii).

All structures show a hexamer consisting of two copies each of IL-11, IL-11Rα and gp130, in agreement with previous immunoprecipitation and native-gel electrophoresis experiments^27^, and a low-resolution EM map of the complex^48^. The structures bear a striking resemblance to a table, with D2 of IL-11Rα and gp130, and IL-11 forming the ‘table-top’, and D3 of IL-11Rα and gp130 forming the ‘legs’ (Figure 1Aii), similar to the IL-6^36^ and vIL-6^35^ complexes. The density for the gp130_EC_ complex shows that D5-D6 of the two gp130 molecules ‘cross over’ (Figure 1Bii), as previously suggested by low-resolution EM reconstructions of the complete extracellular domains of the IL-6^49,50^ and IL-11^48^ signalling complexes, and recent cryoEM studies of the IL-6 signalling complex^51^.

Sedimentation velocity analytical ultracentrifugation (SV-AUC) provided sedimentation coefficient distributions with single, narrow peaks for the gp130_D1-D3_ and gp130_EC_ complexes at 7.1 S and 8.4 S, respectively (Figure 1D). Molecular masses for the gp130_D1-D3_ and gp130_EC_ complexes estimated from SV-AUC of 178.7 kDa (frictional ratio [f/f_0_]: 1.6) and 269.4 kDa (f/f_0_: 1.8) were consistent with masses calculated from sequence (169.8 kDa and 235.0 kDa) as were masses determined by multi-angle light scattering (MALS) of *M*_w_: 187.3 kDa and *M*_w_: 259.6 kDa (Supplementary Figure 3A).

To confirm the solution stoichiometry, we used sedimentation equilibrium analytical ultracentrifugation (SE-AUC), which provides a relative molecular mass (*M*\*) for the complex and its components independent of the extent of glycosylation. SE-AUC of the gp130_D1-D3_ and gp130_EC_ complexes yielded *M*\* of 176.5 kDa and 255.5 kDa respectively (Supplementary Figure 3C-E), in excellent agreement with the sum of the *M\** values obtained for the sum of the hexamer components of 179.2 kDa and 264.4 kDa.

To further probe the solution configuration of the hexamer we collected small-angle X-ray scattering (SAXS) data for the gp130_D1-D3_ and gp130_EC_ complexes (Figure 1E, Supplementary Figure 4A-D, Supplementary Table 3). Our atomic coordinates fit the scattering data well (Figure 1E, Supplementary Table 3), and masses derived from the data of 182.2 kDa and 290.0 kDa agree with calculated hexamer masses. Together these data confirm that our atomic models represent the solution configuration of the hexameric IL-11 signalling complex.

### The IL-11 signalling complex forms in three steps and requires significant structural rearrangement of the cytokine

Our structures of the IL-11 signalling complex allow precise identification of the interactions forming the complex (Figure 2A, B). Overall, complex formation results in the burying of ∼6450 Å^2^ of surface area. The binding sites on IL-11 (Figure 2B) contain residues previously identified by mutagenesis of human and mouse IL-11^52-54^ and by modelling^47^. The structures reveal additional contacts between the loop joining helices A and B of the cytokine (AB loop) and IL-11Rα in site-I and gp130 in site-III that have not previously been identified or interrogated (Figure 2C,D).

**Figure 2:**
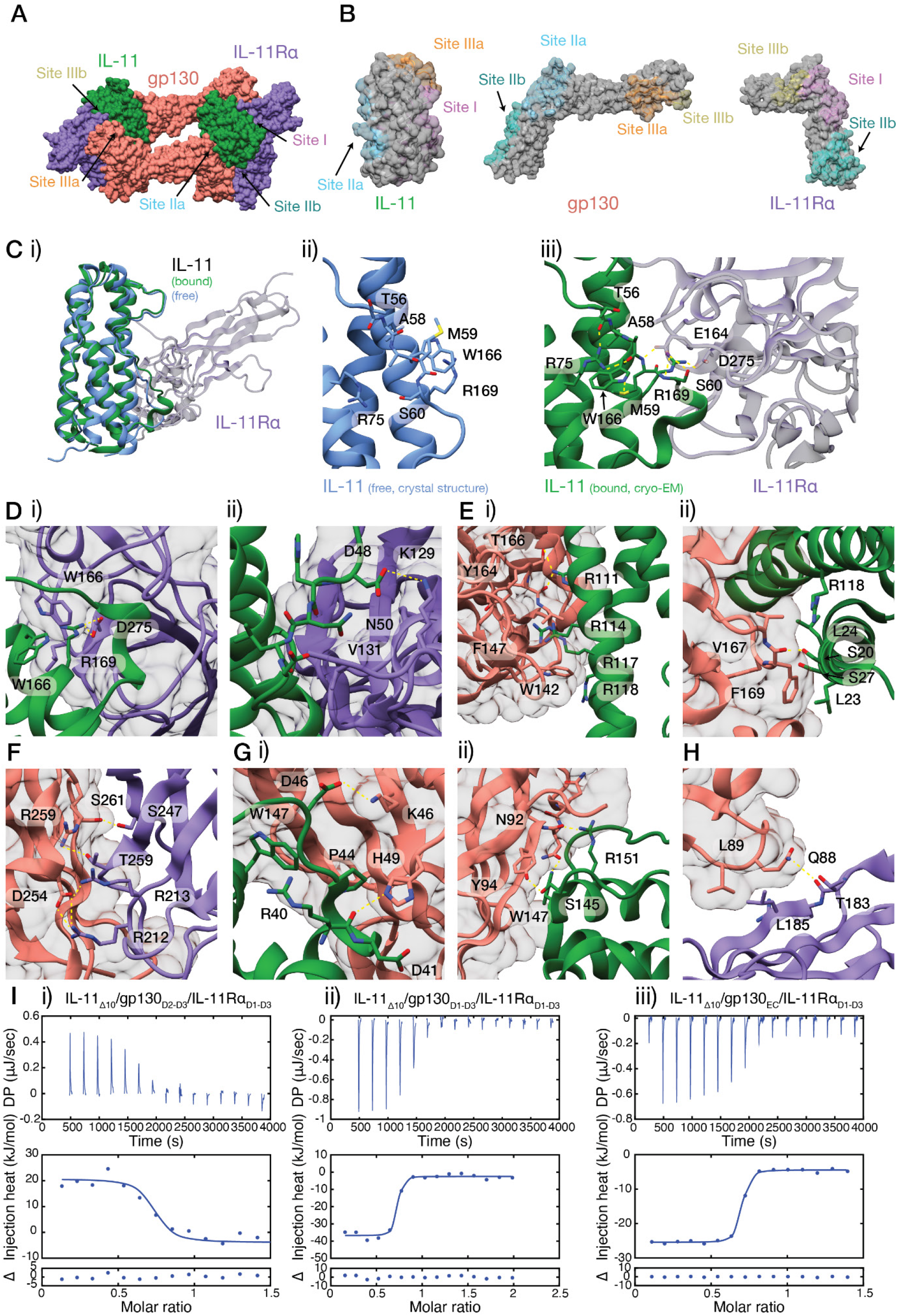
Interactions forming the IL-11 signalling complex. Bound IL-11 is depicted in green, IL-11Rα in purple, and gp130 in salmon. A) Structure of the IL-11_Δ10_/IL-11Rα_D1-D3_/gp130_D1-D3_ complex, with the five binding sites indicated. B) Binding surfaces on IL-11, gp130, and IL-11Rα. C) Rearrangement of the AB loop on complex formation. Uncomplexed IL-11 is depicted in blue. i) overlay of the crystal structure of IL-11^47^ (PDB ID: 6O4O) with the structure of the complex, showing the AB loop rearrangement on complex formation, ii) interactions within the AB loop, and between the AB loop and the α-helical core in the unbound state, iii) the AB loop rearrangement on complex formation. D) Details of site-I contacts (IL-11, green; IL-11Rα, purple; gp130 salmon). i) R169 of IL-11 protrudes into a pocket formed by several hydrophobic residues on IL-11Rα, ii) contacts between the N-terminal end of the AB loop of IL-11 and IL-11Rα. E) Details of site-IIA contacts, i) contacts between C-helix arginine residues of IL-11 and gp130, ii) contacts between the N-terminal end of the A helix of IL-11 and gp130. F) Details of site-IIB contacts. G) Details of site-IIIA contacts, i) contacts between W147 and the N-terminal end of the AB loop of IL-11 with D1 of gp130, ii) contacts between W147 and neighbouring residues of IL-11 with D1 of gp130. H) Details of site-IIIB contacts. D) ITC data for i) the interaction between the IL-11_Δ10_/IL-11Rα_D1-D3_ binary complex and gp130_D2-D3_; ii) the interaction between the IL-11_Δ10_/IL-11Rα_D1-D3_ binary complex and gp130_D1-D3_ and iii), the interaction between the IL-11_Δ10_/IL-11Rα_D1-D3_ binary complex and gp130_EC_. One of three titrations is shown. For complete thermodynamic parameters, see Supplementary Table 4.

Initial formation of a 1:1 complex between IL-11 and IL-11Rα is mediated through site-I of the cytokine (Figure 2C,D). The interaction has an affinity of 23 nM^47^ and is strongly driven by entropy, burying approximately 1000 Å^2^ of surface area. The C-terminal section of the loop joining helices A and B of IL-11 (AB loop; residues F43-G65) undergoes a re-arrangement upon complex formation relative to uncomplexed IL-11^47^ (Figure 2C), resulting in significant conformational change of residues 58-68, including unfolding of the N-terminal turn of helix B. A cation-π interaction between R169 and W166 in uncomplexed IL-11^47^ is broken on complex formation, allowing the AB loop to adopt a new conformation and form extensive contacts with IL-11Rα (Figure 2Cii-iii). The rearrangement allows R169 of IL-11 to protrude into a hydrophobic pocket formed by F165, F230, and L277 of IL-11Rα, and to form a salt bridge with D275. S60 and the backbone amide of M59 within the AB loop both form hydrogen bond networks with E164 of IL-11Rα after rearrangement of the AB loop. The key roles of R169 of IL-11 in mediating both contacts with IL-11Rα and conformational change in the AB loop are consistent with the significant reduction in affinity of IL-11 for IL-11Rα upon substitution of R169 with alanine^47^ (Figure 2Ciii, Figure 2Di). Notably, the new position of the AB loop is stabilised by several new intra-IL-11 contacts, including a hydrogen bond formed by the δ-sulfur of M59 and the indole nitrogen of W166, and two hydrogen bonds formed by the guanidinium guanidium group of R75 with backbone atoms of T56 and A58 in the AB loop (Figure 2Ciii). These conformational changes of IL-11 upon binding IL-11Rα are significantly larger than those observed for other IL-6 family members, IL-6 and LIF^36,38,51^. Several additional contacts are formed by the N-terminal end of the AB loop, including a salt bridge between D48 on the cytokine and K129 on IL-11Rα (Figure 2Dii). However, these contacts do not result in conformational change in this region of the cytokine.

After binding IL-11Rα, IL-11 engages two molecules of gp130, via site-II and site-III, to form the hexameric signalling complex (Figure 2E-H). Site-III interactions are mediated by D1 of gp130 suggesting sequential complex assembly via site-II followed by site-III. To probe the intermediate in IL-11 complex assembly, we generated gp130_D2-D3_, which lacks D1. SV-AUC of the IL-11_Δ10_/IL-11Rα_D1-D3_/gp130_D2-D3_ complex (gp130_D2-D3_ complex) yields a single species with sedimentation coefficient of 4.0 S, and molecular weight of 78.2 kDa (f/f_0_: 1.6; calculated Mw: 73.5 kDa), supporting a trimeric stoichiometry (Supplementary Figure 3F). MALS further supports a trimeric stoichiometry (*M*_w_ = 70.6 kDa, Supplementary Figure 3A). SAXS data for the gp130_D2-D3_ complex fit well to a model of the trimer comprising one molecule of each component assembled via site-I and site-II (Supplementary Figure 4B, E, Supplementary Table 3), further indicating that this trimeric complex is a stable intermediate in the complex assembly.

The interaction between the binary IL-11/IL-11Rα complex and the first molecule of gp130 comprises two coupled interfaces: 1) between IL-11 and gp130 (site-IIA, Figure 2E), and 2) between IL-11Rα and gp130 (site-IIB, Figure 2F). The interactions bury approximately 700 Å^2^ and 600 Å^2^ respectively. The site-IIA interface is hydrophobic in character and the main contact area between IL-11 and gp130 is located on helices A and C of IL-11. Important contacts are mediated by four arginine residues of IL-11 (111, 114, 117 and 118) and two loops of gp130 (residues 142-147 and 164-172) (Figure 2Ei). Key contacts are formed by R114, which protrudes into an aromatic pocket formed by W142 and F147 of gp130. Residues W142 and F169 of gp130 also form important interactions. F169 interacts with the central section of helix A of IL-11, forming hydrophobic contacts with L23 and L24 (Figure 2Eii). F196 of gp130 has been shown to also form key contacts with other IL-6 family cytokines^55,56^. The site-IIB interaction is electrostatic in nature, and results in the formation of ten hydrogen bonds between D3 of IL-11Rα and D3 of gp130 (Figure 2F).

The final interaction to form the complex is a coupled interaction between IL-11 and gp130 D1 (site-IIIA) and between IL-11Rα D2 and gp130 D1 (site-IIIB) (Figure 2G-H). The interaction buries ∼800 Å^2^ in surface area; approximately 600 Å^2^ at site-IIIA, and 200 Å^2^ at site-IIIB. Assembly of the hexamer occurs via two symmetrical site III interfaces between the two IL-11/IL-11Rα/gp130 trimers. The primary contact within site-IIIA is formed by the side chain of W147 of IL-11, which binds flat against a β-strand of gp130 D1 (Figure 2Gi, ii), with the indole nitrogen forming a hydrogen bond to N92 (Figure 2Gi). Contacts are also formed by R151 and S145 of IL-11, which are adjacent to W147 (Figure 2Gi). A second set of interactions are formed by the N-terminal end of the IL-11 AB loop, where P44 packs against D1 of gp130, and D46 forms hydrogen bonds with K46 and N82 of gp130 (Figure 2Gii). A small interface is formed at site-IIIB between a β-sheet (residues 183-187) of IL-11Rα D2 and a loop of gp130 D1 (residues 86-89) (Figure 2H). L185 of IL-11Rα forms hydrophobic contacts with L89 of gp130 and a hydrogen bond is formed between Q88 of gp130 and T183 of IL-11Rα (Figure 2H).

We used isothermal titration calorimetry (ITC) to study the thermodynamics of complex assembly (Figure 2I, Supplementary Table 4). The interaction of the IL-11_Δ10_/IL-11Rα_D1-D3_ binary complex with gp130_D2-D3_ to form the trimer has moderate affinity (*K*_D_ 380 ± 190 nM, ΔH 27 ± 2 kJ/mol, Figure 2Ii), and is strongly driven by entropy. The interaction between the IL-11_Δ10_/IL-11Rα_D1-D3_ complex and gp130_D1-D3_, which includes symmetrical interactions of both molecules of gp130 within the hexamer, is relatively high affinity (*K*_D_ 3 ± 2 nM, ΔH -34± 0.4 kJ/mol, Figure 2Iii). Comparison of the IL-11_Δ10_/IL-11Rα_D1-D3_ binary complex affinities for gp130_D2-D3_ and gp130_D1-D3_ suggest that interaction of a single gp130 molecule via site-III is low affinity and that it is the avidity of the symmetrically duplicated site-III interactions that maintains hexameric complex formation. The affinity of the IL-11Rα_D1-D3_/IL-11_Δ10_ binary complex for gp130_EC_, (*K*_D_ 4 ± 2 nM, ΔH -20 ± 1 kJ/mol, Figure 1Diii) is very similar to the affinity for gp130_D1-D3_, indicating that the membrane-proximal (D4-D6) domains of gp130 do not contribute substantially to complex formation.

Gp130 is shared by most members of the IL-6 family of cytokines. The structure of the hexameric IL-6 signalling complex^36^ showed site-II and site-III interactions with gp130 are similar to those of the IL-11 complex (Supplementary Figure 5A-E), and the LIF/gp130 complex structure comprises a comparable site-II interaction^38^ (Supplementary Figure 5A,B). The three cytokines interact with a similar surface on D2 of gp130 (Supplementary Figure 5A), and IL-6 and IL-11 interact with a similar surface on D1 of gp130, to form the hexameric complex (Supplementary Figure 5D). However, the molecular details of the interactions between the three cytokines differ (Supplementary Figure 5B, C, E, see also Supplementary Discussion).

### The membrane proximal domains of gp130 are highly dynamic

The crystal structure of the extracellular domains of gp130^57^ and low resolution electron microscopy studies of the IL-11^48^, IL-6^49,50^, and LIF^58^ signalling complexes suggest that the membrane proximal domains, D4-D6, are responsible for correctly positioning the transmembrane and intracellular domains of the two signalling receptors. This configuration is thought to orient intracellularly bound JAK molecules for activation, and deletion of any of D4-D6 renders gp130 non-functional *in vitro*^59,60^.

Our consensus maps of the gp130_EC_ complex showed well-defined density for the domains involved in complex formation (D2-D3 of IL-11Rα, IL-11, and D1-D3 of gp130), and poorer density for gp130 D4-D6 (Figure 1B, Supplementary Figure 1Cii, Dii). We considered the possibility that structural dynamics of gp130 D4-D6 contributed to the decreasing density quality with distance from the cytokine binding regions. To explore this, we used 3D variability analysis (3DVA)^61^ in *cryoSPARC*^62^, which resolves conformational changes by fitting a continuous linear subspace model to the cryoEM data (Figure 3, Supplementary Figure 6). 3DVA analysis of the gp130_EC_ complex data reveals three major variability components (Figure 3A-C). The first has minor motion (Figure 3A, Supplementary Movie 1), the second corresponds to a side-to-side oscillation of D4-D6 of gp130 (Figure 3B, Supplementary Movie 2), and the third corresponds to an inward swinging motion of D4-D6 relative to the rest of the complex (Figure 3C, Supplementary Movie 3). Histograms of the distributions along each trajectory are unimodal, indicating continuous motion (Figure 3D, Supplementary Figure 6A-C). This analysis shows that the D4-D6 region of gp130 is dynamic and is not stabilised upon complex formation. The proposed flexibility of D4-D6 is in agreement with our ITC results showing that the gp130_EC_ complex forms with similar affinity to the gp130_D1-D3_ complex, and is supported by our previous molecular dynamics (MD) simulations of gp130 D2-D5 showing that D4 is dynamic with respect to D3^63^. Together these data indicate that D4-D6 of gp130 do not contribute to complex formation. Despite the separation of the N-termini of D6 in the consensus model, the 5-6 residue linker between D6 and the transmembrane helix segment may allow interaction of the two gp130 transmembrane helices of the complex in a parallel configuration (Supplementary Figure 6D). Thus, further studies should examine the D4-D6 domains in the context of the membrane-bound complex, as it is possible that the domains may adopt a more stable conformation in this state.

**Figure 3:**
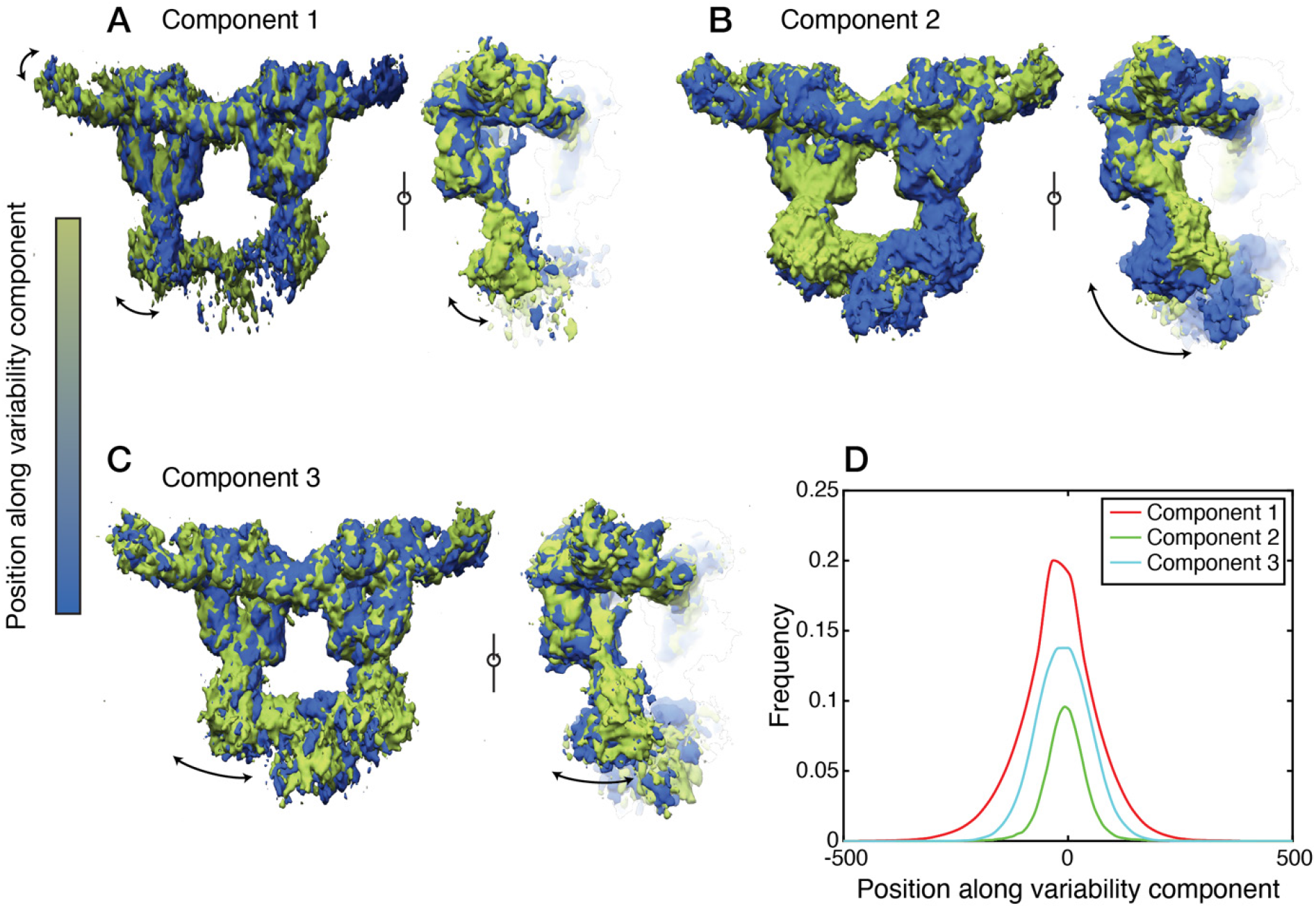
3D variability analysis of the gp130_EC_ complex cryo-EM map. Refined consensus densities along, A) variability component 1, B) variability component 2, C) variability component 3. gp130_EC_ is coloured according to the relative position along the variability component. D) Frequency distributions of the number of particles contributing to each variability component (see also Supplementary Figure 6).

### The cytokine variant “IL-11 Mutein” potently inhibits IL-11 signalling in human cells

IL-11 mutants have previously been proposed to competitively inhibit IL-11 signalling^41,42^. The IL-11 W147A mutation^42^ removes the key tryptophan residue in site-III responsible for the final step of hexameric complex formation (Figure 2Gi). Phage display was subsequently used to identify variants with higher inhibitory potency resulting in “IL-11 Mutein”, which contains W147A combined with mutations of AB loop residues 58-62, ^58^AMSAG^62^ to ^58^PAIDY^62^, that were reported to increase affinity to IL-11Rα^41^.

To characterise these mutants, we generated recombinant human IL-11_Δ10/W147A_, IL-11_Δ10/Mutein_, and a variant containing only the ^58^PAIDY^62^ mutations, which we termed IL-11_Δ10/PAIDY_. We assessed the biological activity of IL-11_Δ10/W147A_, IL-11_Δ10/Mutein_ and IL-11_Δ10/PAIDY_ in Ba/F3 cells stably expressing human gp130 and IL-11Rα^64^ by measuring STAT3 activation, indicated by phosphorylation at Y705 (pSTAT3), using flow cytometry (Figure 4, Supplementary Figure 7). As expected, stimulation with IL-11_Δ10_ produces a robust pSTAT3 response, with an EC_50_ 12 ± 1.1 pM (Figure 4A, Supplementary Figure 7A). Conversely, stimulation with IL-11_Δ10/Mutein_ alone does not result in STAT3 activation, demonstrating that the mutations introduced in IL-11_Δ10/Mutein_ eliminate the ability of IL-11_Δ10/Mutein_ to signal, in agreement with previous reports^41^ (Figure 4A, Supplementary Figure 7A). Surprisingly, stimulation with either IL-11_Δ10/W147A_ or IL-11_Δ10/PAIDY_ resulted in robust STAT3 activation, albeit with a higher EC_50_ than IL-11_Δ10_, of 610 ± 120 pM (p = 0.04 *vs* IL-11_Δ10_) and 140 ± 12 pM (p = 0.01 *vs* IL-11_Δ10_), respectively.

**Figure 4:**
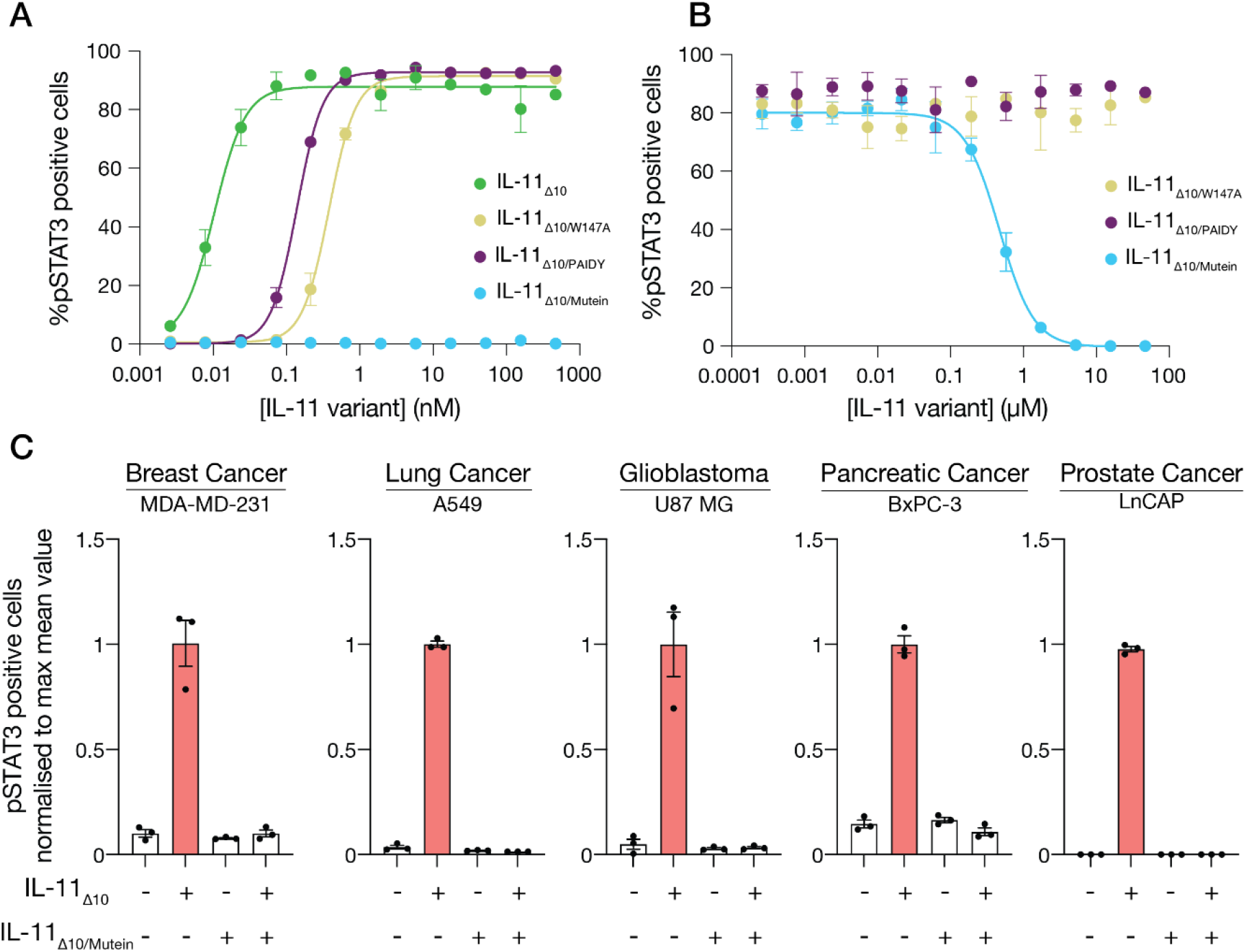
IL-11 Mutein is an effective inhibitor of human IL-11 signalling *in vitro*. A) Representative dose-response curve for IL-11 or IL-11 variant stimulation. The EC_50_ for IL-11_Δ10_ was 0.010 ± 0.002 nM, the EC_50_ for IL-11_Δ10/W147A_ was 0.6 ± 0.2 nM, the EC_50_ for IL-11_Δ10/PAIDY_ was 0.14 ± 0.02 nM, the EC_50_ for IL-11_Δ10/Mutein_ could not be determined. B) Representative dose-response curve for the inhibition of IL-11_Δ10_ stimulation by the IL-11 variants IL-11_Δ10/W147A_, IL-11_Δ10/PAIDY_ and IL-11_Δ10/Mutein_. The IC_50_ for IL-11_Δ10/Mutein_ inhibition was 850 ± 275 nM, the IC50 for the remaining variants was not determined. C) Potent inhibition of IL-11_Δ10_ signalling by IL-11_Δ10/Mutein_ in the indicated human cancer cell lines. Data is presented ± standard error of the mean.

We also examined the inhibitory activity of IL-11_Δ10/Mutein_, IL-11_Δ10/W147A_ and IL-11_Δ10/PAIDY_ in Ba/F3 cells (Figure 4B, Supplementary Figure 7B). IL-11_Δ10/Mutein_ strongly inhibited IL-11_Δ10_ mediated STAT3 activation (IC_50_ = 850 ± 275 nM, Figure 4B), validating IL-11_Δ10/Mutein_ as an effective inhibitor of IL-11 signalling. In contrast, neither IL-11_Δ10/W147A_ or IL-11_Δ10/PAIDY_ inhibited IL-11_Δ10_ mediated STAT3 activation, contradicting previous reports that the W147A mutation alone is an effective IL-11 signalling inhibitor^42,65,66^. Overall, these results show that the inhibitory effect of IL-11_Δ10/Mutein_ requires the combination of both the W147A and ^58^PAIDY^62^ mutations, with either variant alone insufficient to inhibit STAT3 phosphorylation by IL-11.

We extended these studies to several human cell lines and showed that IL-11_Δ10_ robustly stimulates STAT3 activation in the breast cancer cell line MDA-MD-231, the lung cancer cell line A549, the glioblastoma cell line U87-MG, the pancreatic cancer cell line BxPC3, and the prostate cancer cell line LnCap (Figure 4C), in keeping with the tumorigenic function of IL-11. In each of these human cell lines, IL-11_Δ10/Mutein_ effectively inhibits IL-11 mediated STAT3 activation.

### IL-11 Mutein blocks the final step of hexamer assembly

Our functional results prompted investigation of the structural mechanism of IL-11 Mutein signalling inhibition. SV-AUC of the IL-11_Δ10/Mutein_/IL-11Rα_D1-D3_/gp130_D1-D3_ complex yielded a single peak in the sedimentation coefficient distribution of 4.5 S and estimated mass 88.9 kDa (f/f_0_: 1.6), consistent with the formation of a homogenous trimeric complex (Figure 5Ai, Supplementary Figure 8Ai).

**Figure 5:**
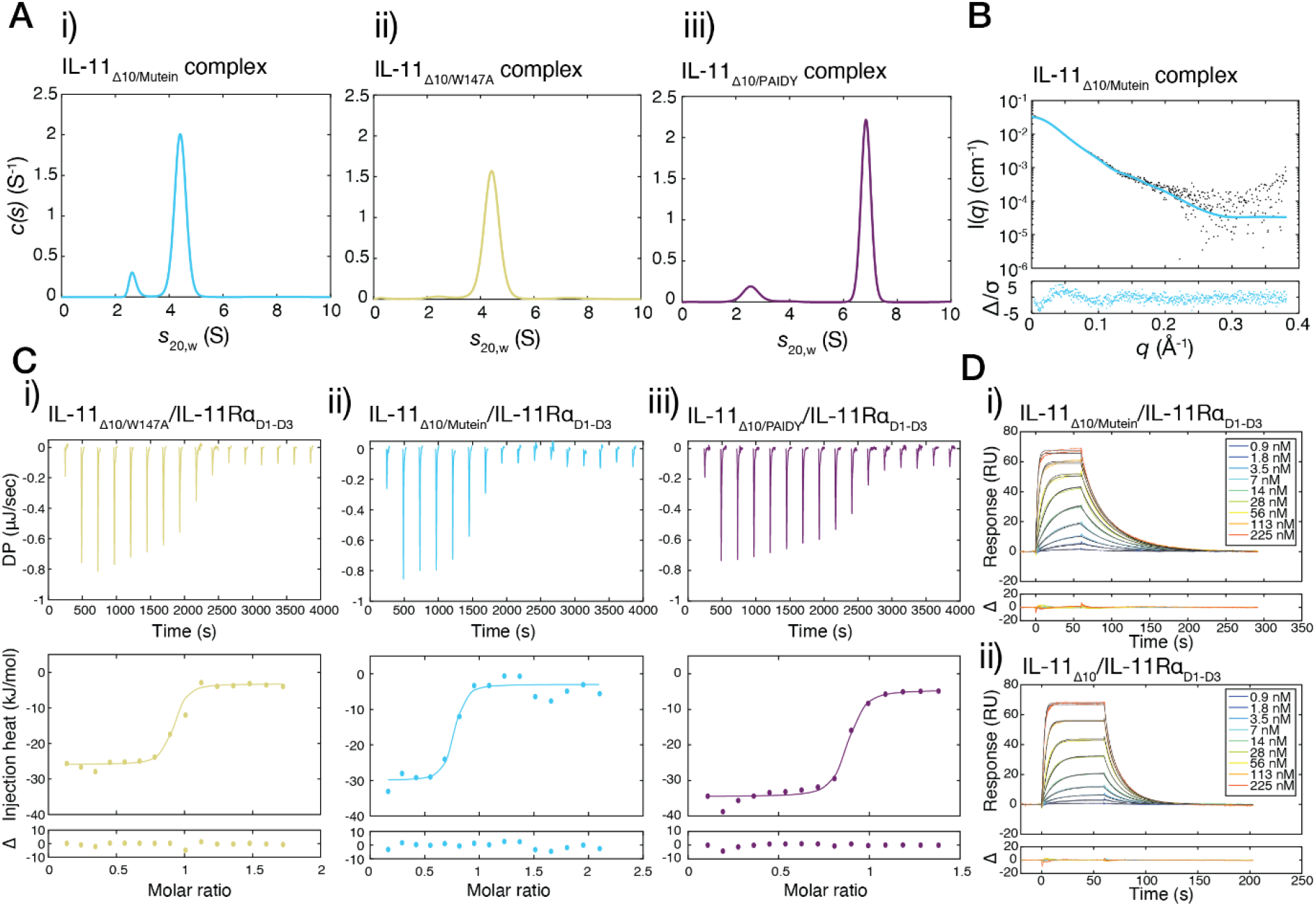
Biophysical basis for IL-11 Mutein signalling inhibition. A) Continuous sedimentation coefficient (c(s)) distributions for the complexes formed between IL-11Rα_D1-D3_, gp130_D1-D3_ and i) IL-11_Δ10_ Mutein, ii) IL-11_Δ10/W147A_, iii) IL-11_Δ10/PAIDY_. B) SAXS data for the IL-11Rα_D1-D3_/gp130_D1-D3_/ IL-11_Δ10_ Mutein complex. The fit shown is to a model of the trimeric complex, χ^2^ 1.7 (see Methods). C) ITC data for the interaction between IL-11Rα_D1-D3_ and i), IL-11_Δ10_ Mutein and ii) IL-11_Δ10/W147A_. D) SPR data for the interaction between IL-11Rα_D1-D3_ and i) biotinylated IL-11_Δ10_ and ii) biotinylated IL-11_Δ10_ Mutein. Black lines show the fit to the data. In both experiments, the biotin tag was used to immobilise IL-11_Δ10_ or IL-11_Δ10_ Mutein to a streptavidin sensor chip. For complete kinetic and thermodynamic parameters for the ITC and SPR experiments, see Supplementary Table 4-5.

SV-AUC of the IL-11_Δ10/W147A_/IL-11Rα_D1-D3_/gp130_D1-D3_ complex indicates that the W147A mutant predominately forms a trimeric complex (sedimentation coefficient: 4.5 S, mass: 82.7 kDa, f/f_0_: 1.5, Figure 5Aii, Supplementary Figure 8Aii). However, a small population of hexameric complex at approximately 7.5 S was observed in the sedimentation coefficient distributions, indicating that the W147A mutation alone permits some hexamer formation and consistent with the ability of IL-11_Δ10/W147A_ to stimulate STAT3 activation *in vitro* (Figure 4). SV-AUC of the IL-11_Δ10/PAIDY_/IL-11Rα_D1-D3_/gp130_D1-D3_ yields a single species with sedimentation coefficient of 6.8 S, and mass of 184.3 kDa (f/f_0_: 1.7), Figure 5Aiii, Supplementary Figure 8Aiii), indicating that IL-11_Δ10/PAIDY_ mediates formation of the hexameric complex to a similar extent to WT IL-11_Δ10_.

Further supporting the trimeric nature of the IL-11_Δ10/Mutein_ complex, SAXS data collected on the IL-11_Δ10/Mutein_/gp130_D1-D3_/IL-11Rα_D1-D3_ complex (Figure 5B, Supplementary Figure 8B) agrees well with the calculated scattering profile of a single trimer of the gp130_D1-D3_ complex. Furthermore, *ab initio* models from SAXS data collected on the IL-11_Δ10/Mutein_/IL-11Rα_D1-D3_/gp130_D1-D3_ complex and the gp130_D1-D3_ complex hexamer are consistent with the IL-11_Δ10/Mutein_ complex being half the size in one dimension compared to the hexameric complex (Supplementary Figure 9C). The mass of the IL-11_Δ10/Mutein_ complex determined by SV-AUC and SAXS is also supported by the mass determined using MALS (*M*_w_ 79.6 kDa, Supplementary Figure 9D).

In combination, these results indicate that that both IL-11_Δ10/W147A_ and IL-11_Δ10/PAIDY_ can mediate assembly of the hexameric signalling complex, albeit to differing extents, while formation of site III interactions by IL-11_Δ10/Mutein_ is abrogated, and assembly is stalled at the trimer stage.

### IL-11 Mutein/IL-11Rα binding kinetics are different to WT IL-11

The ^58^PAIDY^62^ mutations present in the AB loop of IL-11 were proposed to increase the affinity for IL-11Rα twenty-fold over WT IL-11, resulting in effective competition for IL-11Rα binding^41^. To test this, we measured the affinity of the mutant cytokines for IL-11Rα using ITC (Figure 5C). IL-11_Δ10/W147A_, IL-11_Δ10/Mutein_, and IL-11_Δ10/PAIDY_ interact with IL-11Rα_D1-D3_ with *K*_D_ of 10 ± 8 nM, 38 ± 9 nM, (p = 0.2 *vs* IL-11_Δ10/W147A_), and 81 ± 44 nM, (p = 0.3 *vs* IL-11_Δ10/W147A_) respectively (Figure 5C). These data suggest that the PAIDY mutations do not significantly increase the affinity of the IL-11 for IL-11Rα.

We also measured binding kinetic parameters for IL-11_Δ10_ and IL-11_Δ10/Mutein_ with IL-11Rα_D1-D3_ using surface plasmon resonance (SPR). IL-11 constructs were C-terminally biotinylated *via* an avitag^67^ and coupled to a streptavidin SPR chip (Figure 5D). Despite similar affinity, the kinetics of the interactions are different, with approximately three-fold slower dissociation rate for IL-11_Δ10/Mutein_/IL-11Rα_D1-D3_ (*k*_d_ 0.34 ± 0.01 × 10^−1^ s^-1^, Figure 5Di) than IL-11_Δ10_/IL-11Rα_D1-D3_ (*k*_d_ 1.1 ± 0.2 × 10^−1^ s^-1^, Figure 5Dii n=2; Supplementary Table 5; p = 0.06 *vs* IL-11_Δ10/Mutein_). These results show that the PAIDY mutations present in IL-11_Δ10/Mutein_ alter the kinetics of interaction with IL-11Rα, compared to IL-11, but do not significantly alter the affinity, suggesting that the mechanism of signalling inhibition by IL-11 Mutein is more complex than previously appreciated.

### The conformation of the AB loop of IL-11 Mutein is altered relative to WT IL-11

We solved crystal structures of IL-11_Δ10/Mutein_ and IL-11_Δ10/W147A_ at resolutions of 1.8 Å and 1.5 Å, respectively, to understand the structural consequences of the mutations (Figure 6A-B, Supplementary Figure 9A-B, Supplementary Table 2). SAXS experiments show that the crystal structures are representative of the solution structure, and SV-AUC indicates that both proteins are monomeric in solution (Supplementary Figure 9C-D, Supplementary Figure 10).

**Figure 6:**
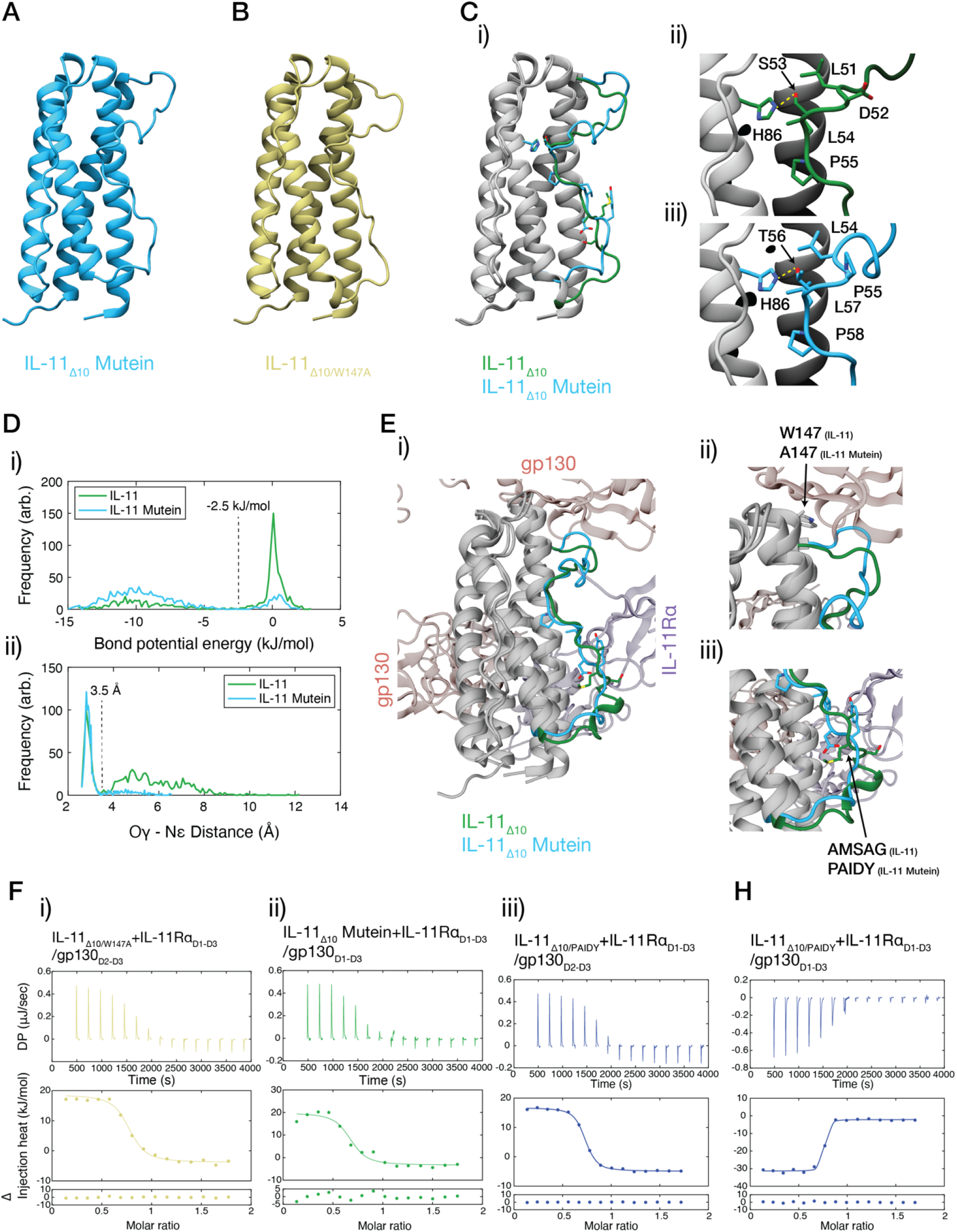
The structure of IL-11 Mutein, and mechanism of inhibition. A) The structure of IL-11_Δ10_ Mutein. B) The structure of IL-11_Δ10/W147A_. C) Overlay of IL-11_Δ10_ Mutein and IL-11_Δ10_ (PDB ID: 6O4O^47^), i) overall view of both structures, with the AB loop coloured as indicated in the figure, ii) detail of the AB loop, showing changes in loop-core interaction that occur as a result of the PAIDY mutations in IL-11 Mutein. D) MD analysis of the hydrogen bond between S53/T56 and H86, i) distribution of the estimated bond potential energy for IL-11Δ10 and IL-11Δ10 Mutein through the simulation, an approximate cut-off for hydrogen bonding is indicated, ii) distance distribution for the distance between the donor oxygen (S53/T56 Oγ) and the acceptor nitrogen (H86 Nε) through the simulation, an approximate cut-off for hydrogen bonding is indicated. E) Overlay of the structure of IL-11_Δ10_ Mutein with the structure of the IL-11Rα_D1-D3_/gp130_D1-D3_/ IL-11_Δ10_ complex, i) overall view, ii) detail of the site-III interface, with the W147/A147 residue shown, iii) detail of the AB loop, with the AMSAG/PAIDY residues shown. F) ITC data for the interaction between IL-11_Δ10_ Mutein/IL-11Rα_D1-D3_ and gp130_D1-D3_. G) ITC data for the interaction between IL-11_Δ10/W147A_/IL-11Rα_D1-D3_ and gp130_D1-D3_. H) ITC data for the interaction between IL-11_Δ10/W147A_/IL-11Rα_D1-D3_ with i) gp130_D2-D3_ and ii) gp130_D1-D3_. For complete thermodynamic parameters, see Supplementary Table 4, for ITC experiments one of three representative titrations are shown.

The structure of IL-11_Δ10/W147A_ is similar to the structure of IL-11_Δ10_ (Supplementary Figure 9Ei, RMSD 0.3 Å relative to IL-11_Δ10_^47^), indicating that the W147A mutation does not alter the structure. In contrast, the AB loop of IL-11_Δ10/Mutein_ shows significant conformational shifts relative to the WT structure caused by alterations in key contacts between the loop and the four-α-helical bundle (Figure 6Ci, Supplementary Figure 9Eii, RMSD 1.3 Å relative to IL-11_Δ10_^47^).

In IL-11_Δ10_^47^ and IL-11_Δ10/W147A_, a hydrogen bond is formed between the side chains of S53 within the AB loop and H86 on the helical bundle. In contrast, in IL-11_Δ10/Mutein_ this hydrogen bond is formed between T56 of the loop and H86 (Figure 6C, for overlay see Supplementary Figure 9F). Similarly, key hydrophobic interactions between AB loop residues L51, L54 and P55 and the helical core structure of IL-11_Δ10_ and IL-11_Δ10/W147A_ (Figure 6Cii, Supplementary Figure 9Gi) are mediated by L54, L57 and P58 in IL-11_Δ10/Mutein_ (Figure 6Ciii, Supplementary Figure 9Gii). Thus, the ^58^PAIDY^62^ mutations alter the position of the AB-loop, shifting the register of key loop-core interactions by three residues, while maintaining the composition of the interacting residues. This register change is due to a shift of the ‘LX(S/T)LP’ motif that mediates loop-core contacts: In IL-11_Δ10_ and IL-11_Δ10/W147A_ the motif is ^51^LDSLP^55^, while in IL-11_Δ10/Mutein_ the interacting motif is ^54^LPTLP^58^. The result is that the bulk of the loop is shifted toward site-III and the N-terminal turn of helix-B is unfolded. We note that this configuration of the C-terminal segment of the AB-loop and N-terminal turn of helix-B resembles the structure of wild-type IL-11_Δ10_ bound within the hexameric complex.

To assess the dynamics of the AB loop, we performed 1 μs molecular dynamics simulations of both IL-11_Δ10_ and IL-11_Δ10/Mutein_ (Figure 6D, Supplementary Figure 11, Supplementary Movie 4-5). The overall dynamic profile of the four-α-helical core of the two proteins is very similar (Supplementary Figure 11A-C). We analysed the stability of the IL-11_Δ10_ S53-H86 hydrogen bond and the IL-11_Δ10/Mutein_ T56-H86 hydrogen bond throughout the simulation by calculating the distance between the donor (S53 Oγ or T56 Oγ) and the acceptor (H86 Nε), and the distribution of the potential hydrogen bond energy (Figure 6D, Supplementary Figure 11D-F). Using a hydrogen bond energy cut-off of -2.5 kJ/mol, S53 of IL-11_Δ10_ is hydrogen bonded to H86 for 33% of the simulation, while T56 of IL-11_Δ10/Mutein_ is hydrogen bonded to H86 for 84% of the trajectory. This increased persistence of the hydrogen bond suggests that the AB loop conformation of IL-11_Δ10/Mutein_ is more stable than that of IL-11_Δ10_. Differential scanning fluorometry (DSF) analysis revealed that the temperature of hydrophobic exposure (*T*_h_^68^) of IL-11_Δ10/Mutein_ was 88.7 ± 0.16 °C (mean ± SE, n = 3) indicating significantly higher thermal stability than both IL-11_Δ10_ (*T*_h_ 84.8 ± 0.39 °C, p = 0.003 *vs* IL-11_Δ10/Mutein_) and IL-11_Δ10/W147A_ (*T*_h_ 87.3 ± 0.29 °C, p = 0.04 *vs* IL-11_Δ10/Mutein_) (Supplementary Figure 12A).

Together these data indicate that altered loop-core interactions brought about by the ^58^PAIDY^62^ mutations in IL-11_Δ10/Mutein_ result in a new, stabilised position of the AB loop. We propose that this improved stability is due to the backbone structural restraints introduced in the IL-11_Δ10/Mutein_ LX(S/T)LP motif by P55 at the ‘X’ position.

### The AB loop of IL-11 Mutein acts in combination with the W147A mutation to block hexamer formation at site-III

Our IL-11 complex structures show that the N-terminal section of the AB loop contacts gp130 at site-III, suggesting that the altered conformation and stability of the AB loop in IL-11_Δ10/Mutein_ could alter interactions at this site. Superposition of the crystal structure of IL-11_Δ10/Mutein_ with an IL-11 molecule of our gp130_D1-D3_ complex structure (Figure 6E) revealed that the N-terminal part of the AB-loop clashes sterically with D1 of gp130 at site-III in the complex (Figure 6Eii). This observation suggested that the altered conformation of the AB-loop acts to disrupt gp130 binding at site-III.

To investigate this possibility, we determined the affinities of the IL-11_Δ10/W147A_/IL-11Rα_D1-D3_, IL-11_Δ10/PAIDY_/IL-11Rα_D1-D3_, and IL-11_Δ10/Mutein_/IL-11Rα_D1-D3_ binary complexes for gp130 (Figure 6F-G, Supplementary Table 4). The IL-11_Δ10/W147A_/IL-11Rα_D1-D3_ binary complex binds gp130_D2-D3_, forming a trimeric complex, with similar affinity to IL-11_Δ10_ (*K*_D_ 130 ± 14 nM, p = 0.15 *vs* IL-11_Δ10_, Figure 6Fi). In contrast, the interaction of IL-11_Δ10/Mutein_/IL-11Rα_D1-D3_ with gp130_D1-D3_ and IL-11_Δ10/PAIDY_/IL-11Rα_D1-D3_ with gp130_D2-D3_ to form a trimer was significantly higher affinity than WT IL-11_Δ10_, with *K*_D_ of 55 ± 4 nM, (p = 0.01 *vs* IL-11_Δ10_) (Figure 6Fii) and 60 ± 16 nM (p = 0.04 *vs* IL-11_Δ10_) (Figure 6Fiii), respectively. Interestingly, IL-11_Δ10/PAIDY_/IL-11Rα_D1-D3_ binds gp130_D1-D3_ to form the hexameric complex with similar affinity compared to IL-11_Δ10_ (*K*_D_ 20 ± 17 nM, p = 0.2 *vs* IL-11_Δ10_, Figure 6H, Supplementary Table 4).

These ITC results show that the PAIDY mutations increase site-II affinity but are insufficient to disrupt hexamer formation in the absence of the W147A mutation These observations are consistent with our biological assays (Figure 4), indicating that the synergistic effects of the PAIDY and W147A mutations are required for the potent antagonist function of IL-11 Mutein.

The increase in site-II affinity of IL-11_Δ10/Mutein_ and IL-11_Δ10/PAIDY_ was surprising, given that the mutations in both proteins lie distant from the site-II interface (Figure 6E). This increased affinity may be explained by the decreased dissociation rate of the IL-11_Δ10/Mutein_/IL-11Rα_D1-D3_ complex relative to the IL-11_Δ10_/IL-11Rα_D1-D3_ complex. In this case, the increased residence time of IL-11_Δ10/Mutein_ bound to IL-11Rα_D1-D3_ increases the period that the composite site-II binding interface of gp130 is intact and, therefore, increases the microscopic association rate of gp130 while having little effect on its dissociation rate (Figure 5E). Alternatively, the altered site-II interaction may reflect modified dynamics of the bound cytokine variant or subtle adjustment of its binding pose on IL-11Rα induced by the ^58^PAIDY^62^ mutations.

Our ITC and SPR results, in combination with the structures of the IL-11 signalling complex, IL-11_Δ10/Mutein_, and IL-11_Δ10/W147A_ show that the ^58^PAIDY^62^ mutations in IL-11 Mutein significantly alter the interaction of the cytokine with both IL-11Rα and gp130 at all three binding sites, which likely underpins the efficacy of IL-11 Mutein as an IL-11 signalling inhibitor. The slower dissociation rate at site I and increased affinity at site II enhance the ability of IL-11_Δ10/Mutein_ to compete with native IL-11 for binding to IL-11Rα and gp130, thereby contributing to the competitive inhibition mechanism. These results also show that the AB loop in IL-11 is a critical region for the formation of the signalling complex, which will guide the design of future inhibitors.

## Discussion

IL-11 signalling has been implicated in a number of human diseases and is pre-clinically validated as a therapeutic target, prompting significant investment in therapeutic development, with the first clinical trials imminent. However, informed development of IL-11 signalling inhibitors has been hindered by a lack of structural knowledge of the IL-11 complex, and a lack of understanding of the mechanism of action of existing IL-11 signalling inhibitors. We report the first structures of the IL-11 signalling complex, including information on the position and dynamics of the extracellular domains of gp130, which have previously eluded high-resolution structural characterisation. We show that the complex forms in three steps involving significant conformational rearrangement of the cytokine, and that the nature of the interactions forming the complex is distinct when compared to the other well-characterised IL-6 family cytokines, IL-6 and LIF.

The formation of the signalling complex results in activation of JAKs bound to the cytoplasmic region of gp130, mainly JAK1 and to a lesser extent JAK2 and TYK2s. Activated JAKs subsequently phosphorylate tyrosine residues of gp130 resulting in activation of signalling pathways, including STAT, ERK, MAPK and phosphoinositide 3-kinase (PI3K) pathways that mediate biological outcomes^29,30^. Our data on the nature and dynamics of the membrane proximal domains of gp130 have implications for understanding the molecular mechanisms of JAK activation and propagation of downstream signalling pathways and will inform further interrogation of signalling complexes including transmembrane and cytoplasmic machinery. In future, this knowledge may inform how these pathways are co-activated, or individually activated in disease, as evidence emerges that there may be a cell context dependent IL-11 mediated activation of STAT3 pro-survival pathways compared to IL-11 mediated ERK driven proliferation and apoptosis in disease^69^. Importantly, by defining the structural and biophysical dynamics of gp130 within the signalling complex our data enable a better understanding of the pathogenicity of emerging cytokine selective variants in *IL6ST*^63,70-75^, as well as variants in *IL11R*^76,77^ and *IL11*^78,79^, for which there are no targeted therapeutic opportunities for patients.

Our characterisation of the IL-11 signalling inhibitor, IL-11 Mutein, shows that it is highly effective at blocking signalling complex formation and potently inhibits signalling in a range of human cancer cell lines, highlighting the breadth of disease applications for emerging IL-11 signalling therapeutics. In contrast, the point mutant, IL-11 W147A, does not antagonise signalling by wild-type IL-11, contradicting previous reports^42,65,66^. In particular, our data show the importance of the IL-11 Mutein AB loop in modulating interactions with IL-11Rα and both gp130 molecules of the complex leading to inhibition of signalling. Thus, engineering of the AB loop may be a general path for generation of antagonist variants of class I cytokines. Our results functionally and mechanistically validate IL-11 Mutein as a tool IL-11 signalling inhibitor and provide a facile, high-yield method for producing IL-11 Mutein suitable for laboratory studies of IL-11 biology.

Overall, the results presented here will facilitate significant advancement of IL-11 biology and will appreciably aid the development of therapeutic agents targeting IL-11 and related cytokines.

## Acknowledgements

Parts of this research were conducted at the SAXS/WAXS and MX2 beamlines of the Australian Synchrotron, part of the Australian Nuclear Science and Technology Organisation, and made use of the ACRF Detector at the MX2 beamline. Initial crystallisation screens were conducted at the CSIRO Collaborative Crystallisation Centre, Melbourne, Australia. This research made use of the ACRF Facility for Innovative Cancer Drug Discovery and the LIEF HPC-GPGPU Facility hosted at the University of Melbourne (LE170100200). We acknowledge the Ian Holmes Imaging Centre, the Melbourne Protein Characterisation Platform, and the Melbourne Mass Spectrometry and Proteomics Facility, at the Bio21 Molecular Science and Biotechnology Institute, for exceptional technical support.

This work was supported by the National Health & Medical Research Council of Australia (APP1147621, APP1080498). R.D.M. was the recipient of an Australian Government Research Training Program Scholarship. T.L.P. is the recipient of a Sylvia and Charles Viertel Senior Medical Research Fellowship. Funding from the Victorian State Government Operational Infrastructure Support Scheme is acknowledged.

## Author Contributions

R.D.M., E.H., T.L.P., and M.D.W.G. conceived the studies, designed and performed experiments, analysed data, and wrote the manuscript. K.Y.F., K.A., C.O.Z., C.C.K., L.D., C.M., and A.P.L. designed and performed experiments, and analysed data. M.W.P. P.R.G contributed critical resources and intellectual input. All authors have read and agreed to the manuscript content.

## RESOURCE AVAILABILITY

### Lead Contact

Correspondence should be direct to the lead contact, Associate Professor Michael Griffin.

## Materials availability

Unique materials generated in this study may be made available to qualified, academic, non-commercial researchers upon request to the lead contact with a completed materials transfer agreement.

## Data availability

Structure coordinates, cryo-EM density maps, and structure factors have been deposited in the Protein Data Bank and Electron Microscopy Data Bank. All other data are contained within the manuscript and Supplemental Information.

## METHOD DETAILS

### Protein expression and purification

IL-11, IL-11 mutants, and IL-11Rα constructs were expressed and purified as previously described^47^. Gp130 constructs were expressed using the same insect cell expression method as IL-11Rα^47^. Throughout this work, we used an N-terminally truncated form of IL-11 (IL-11_Δ10_), and a C-terminally truncated form of IL-11Rα (IL-11Rα_D1-D3_), containing the structured extracellular domains characterised previously^47^. We used gp130 constructs containing the D1-D3 domains of gp130 (gp130_D1-D3_), D2-D3 domains (gp130_D2-D3_), or D1-D6 domains of gp130, which is the complete extracellular region of gp130 (gp130_EC_)^57^.

For coupling to a streptavidin SPR chip, we generated IL-11_Δ10_ and IL-11_Δ10/Mutein_ constructs with a C-terminal avitag^67^, which was biotinylated *in vitro* in *Escherichia coli* BL21(DE3) cells. These fusion proteins were purified using identical methods to IL-11, and the degree of biotinylation was assessed using mass spectrometry.

### Purification of the IL-11 signalling complex

The IL-11 signalling complex was prepared by mixing equimolar amounts of IL-11_Δ10_, IL-11Rα_D1-D3_ and gp130_D1-D3_/gp130_EC_. The complex was incubated on ice for approximately one hour, then applied to a Superdex 200 10/30 size exclusion column, pre-equilibrated in TBS pH 8. Fractions containing the signalling complex were pooled and concentrated to 1-5 mg/mL. Purity of the complex was assessed using native-PAGE electrophoresis and sedimentation-velocity analytical ultracentrifugation. The IL-11_Δ10_/IL-11Rα_D1-D3_/gp130_EC_ complex was prepared using an identical method, although the complex used for cryoEM was not size exclusion purified.

### Cryo-electron microscopy – data collection and 3D reconstruction

Cryo-EM was performed at the Bio21 Institute Ian Holmes Imaging Centre. The IL-11_Δ10_/IL-11Rα_D1-D3_/gp130_D1-D3_ complex (0.5 mg/mL) was blotted onto UltrAuFoil grids (R2/2, Quantifoil Micro Tools GmbH) that had been subjected to glow discharge (15 mA for 30 s). Grids were imaged using a Gatan K2 direct detector mounted on a Talos Arctica (FEI, Hillsborough, Oregon) with a 70-µm objective aperture. The detector was operated in super-resolution counting mode at 0.655 Å/ pixel (100,000 × magnification) with a defocus range of -0.8 to - 2.0 µm. 40 frames per movie were acquired for a total dose of 50 electrons Å^-2^. Movies were acquired with specimen grids both untilted and with tilts of up to 35°.

The IL-11_Δ10_/IL-11Rα_D1-D3_/gp130_EC_ complex (1.5 mg/mL) in the presence of 1-2 mM *n*-dodecyl β-D-maltoside was blotted onto UltrAuFoil grids (R2/2, Quantifoil Micro Tools GmbH) that had been subjected to glow discharge (15 mA for 30 s). Grids were imaged using a Gatan K2 direct detector mounted on a Talos Arctica (FEI, Hillsborough, Oregon) with a 70-µm objective aperture. The detector was operated in counting mode. The complex was imaged at 1.31 Å/ pixel (100,000 × microscope magnification) with a defocus range of -0.8 to -2.0 µm. 40 frames per movie were acquired for a total dose of 50 electrons Å^-2^.

Analyses of the IL-11_Δ10_/IL-11Rα_D1-D3_/gp130_D1-D3_ complex were carried out using RELION-3.0^80,81^. Movie motion was corrected using MotionCor2.1. Cryosparc 2.1^62^ was used for CTF estimation using the patch CTF estimation routine. A total of 3,082,536 particles were extracted from 2,010 motion-corrected movies. After 2D class averaging in Cryosparc 2.1 625,866 particles were retained and were re-extracted in RELION-3.0 with the original defocus from Cryosparc 2.1. After 3D classification 204,455 particles were used for the final 3D refinement. Final refinement with C2 symmetry yielded a map with a resolution of 3.5 Å. Resolution was estimated using gold standard FSC = 0.143 calculated using a relaxed solvent map. Maps were sharpened using *phenix*.*auto_sharpen*^82^. Local resolution maps were calculated using *Resmap*^83^. Buried surface area was determined using *PISA*.

Analysis of the IL-11_Δ10_/IL-11Rα_D1-D3_/gp130_EC_ complex was carried out using cryosparc 2.1 Movie motion was corrected using MotionCor2.1. Cryosparc 2.1^62^ was used for CTF estimation using the patch CTF estimation routine. WARP^84^ was used to pick 694,360 particles from 6,861 motion corrected movies. After 3 rounds of 2D classification, particles were submitted to heterogenous refinement and one further 2D classification leading to a final particle number of 125,373. These particles were used for standard refinement (3.99 Å) followed by a final non-uniform refinement using C2 symmetry leading to a map with a resolution of 3.76 Å.

### Cryo-electron microscopy – Model building and refinement

For the IL-11_Δ10_/IL-11Rα_D1-D3_/gp130_D1-D3_ complex, published models of IL-11^47^ (PDB ID: 6O4O), gp130 D1-D3 (chain A from PDB ID: 1I1R^35^), and the IL-11Rα chain from an antibody-bound structure of IL-11Rα were docked into the cryoEM density map of the IL-11_Δ10_/IL-11Rα_D1-D3_/gp130_D1-D3_ complex using *UCSF Chimera* to generate an initial model of the complex. This model was refined using *phenix*.*real_space_refine*^85-87^, followed by manual model-building in *Coot*^85^ and automated refinement in *phenix*.*real_space_refine*. Strict NCS was enforced in refinement. Reference model restraints were used throughout refinement, using the initial models as the reference models. Geometry validation was performed using the *phenix*.*validation_cryoem* tool (incorporating *MOLProbity*) and *EMRinger*^88,89^. Structures were visualised using *UCSF Chimera*^87^. Structures were aligned using the *Matchmaker* algorithm in *UCSF Chimera* or the *CE* algorithm^90^ in *PyMOL* 2.2. Map-model FSC curves were calculated in *Phenix*. Figures were prepared using *UCSF Chimera*^87^.

For the IL-11_Δ10_/IL-11Rα_D1-D3_/gp130_EC_ complex, published models of IL-11^47^ (PDB ID: 6O4O), gp130 D1-D3 (chain A from PDB ID: 1I1R^35^) gp130_D4-D6_^57^ (PDB ID: 3L5I), and the IL-11Rα chain from an antibody-bound structure of IL-11Rα were docked into the cryoEM density map of the IL-11_Δ10_/IL-11Rα_D1-D3_/gp130_D1-D3_ complex using *UCSF Chimera* to generate an initial model of the complex. Refinement was conducted in the same manner as the IL-11_Δ10_/IL-11Rα_D1-D3_/gp130_D1-D3_ complex. The deposited model of the IL-11_Δ10_/IL-11Rα_D1-D3_/gp130_EC_ complex did not include the D5-D6 domains of gp130; we prepared a second model, which we did not deposit, which included the D5-D6 domains of gp130.

The conclusions regarding the binding interactions described are supported by the gp130_D1-D3_ and gp130_EC_ complex structures, and the crystal structure of the complex. Figure 2 was prepared with the gp130_D1-D3_ complex structure, as this structure is supported by the highest resolution data.

For the IL-11_Δ10_/IL-11Rα_D1-D3_/gp130_EC_ complex 3D variability analysis with 3 modes to solve was performed in C1 symmetry on the final 125,373 particles used for refinement without symmetry expansion. 3D variability display was used in intermediate mode with 9 frames for each mode solved.

### Protein crystallisation and X-ray data collection

The IL-11 signalling complex for crystallisation was prepared by combining equimolar amounts of IL-11_FL_, IL-11Rα_EC_ and gp130_D1-D3_, followed by purification of the complex using gel filtration. Crystals of the IL-11 signalling complex were grown at 20 °C in the condition 180 mM magnesium chloride, 15.3% PEG 3350, 100 mM potassium sodium tartrate, 90 mM sodium HEPES pH 7.25 and 1.8% tert-butanol. Crystallisation drops were prepared by mixing equal volumes of the precipitant and IL-11 signalling complex (at 3 mg/mL), and 10 μL of a seed stock (prepared according to the method of Luft and DeTitta^91^). Crystals were flash-cooled in liquid nitrogen directly from crystallisation drops, and X-ray diffraction data were collected at 100 K at the Australian Synchrotron MX2 beamline^92^. X-ray data collection statistics are tabulated in Supplementary Table 2.

Initial crystals of IL-11_Δ10/Mutein_ were obtained using a similar screening approach to IL-11_Δ1047_, in the precipitant 30% PEG 3350, 0.2 M ammonium sulfate, 0.1 M Tris pH 8.5, 20 °C. These initial crystals were used to prepare a microseed stock^91^. Large, single plates of IL-11_Δ10/Mutein_ grew in the condition 27% PEG 3350, 0.1 M bis-tris propane pH 9, 0.2 M ammonium sulfate, 5 mM praseodymium chloride, 20 °C. Crystallisation drops were produced by mixing 1.5 μL precipitant, 1.5 μL IL-11_Δ10/Mutein_ (5 mg/mL) and 0.5 μL seed. Crystals of IL-11_Δ10/Mutein_ were large plates, with approximate dimensions 200 × 200 × 5 μm. A similar approach was used to crystallise IL-11_Δ10/W147A_. These mutants were cross seeded with seed generated from IL-11_Δ10_ or IL-11_Δ10/Mutein_ crystals and grew in very similar conditions to IL-11_Δ10/Mutein_. Crystals of IL-11_Δ10/W147A_ were rods, similar to crystals of IL-11_Δ10_^47^.

Crystals were flash-cooled in liquid nitrogen directly from crystallisation drops, and X-ray diffraction data were collected at 100 K at the Australian Synchrotron MX2 beamline^92^. X-ray data collection statistics are tabulated in Supplementary Table 2.

### X-ray data processing and structure refinement

For the IL-11 signalling complex, diffraction data were indexed, integrated and scaled using *XDS*^93^, analysed using *POINTLESS*^94^ and merged using *AIMLESS*^95^ from the *CCP4* suite. Due to the highly anisotropic nature of the data, an ellipsoidal resolution cutoff was applied using the *STARANISO*^96^ server. Initial phase estimates were obtained using molecular replacement with *Phaser*^97^ using the cryoEM structure of the IL-11_Δ10_/IL-11Rα_D1-D3_/gp130_D1-D3_ complex as the search model. Refinement was performed in *phenix*.*refine*^98^ with rigid-body refinement. Strict NCS restraints (NCS constraints) were used in refinement. Reference model restraints and Ramachandran restraints were used, using the high-resolution structures of the complex components. Iterative model-building was performed using *Coot*^85^ using NCS-averaged maps. NCS map averaging was performed in *Coot*. Refinement statistics are tabulated in Supplementary Table 2.

For IL-11_Δ10/Mutein_, and IL-11_Δ10/W147A_, diffraction data were indexed, integrated and scaled using *XDS*^93^, analysed using *POINTLESS*^94^ and merged using *AIMLESS*^95^, initial phase estimates were obtained using molecular replacement with *Phaser*^97^, using either our original structure of IL-11 (PDB ID: 4MHL)^79^ for IL-11_Δ10/Mutein_ or our high-resolution structure of IL-11_Δ10_ for IL-11_Δ10/W147A_ (PDB ID: 6O4O)^47^ as the search model. Auto-building with simulated annealing was performed in *phenix*.*autobuild* to reduce phase bias from the search model. Refinement was performed in *phenix*.*refine*^98^ with iterative manual building using *Coot*^*85*^. TLS refinement was performed using a single TLS group containing all protein atoms. Explicit riding hydrogens were used throughout refinement and included in the final model, the atomic position and *B* factors for hydrogens were not refined. Residues of all structures are numbered in an identical manner to our structure of IL-11_Δ10_ (PDB ID: 6O4O)^47^, reflecting the mature protein sequence after cleavage of signal peptide. Structures were aligned using the CE^90^ algorithm in *PyMOL* 2.2. Refinement statistics are tabulated in Supplementary Table 2.

### Analytical ultracentrifugation

SV-AUC experiments were conducted using a Beckman Coulter XL-I analytical ultracentrifuge or a Beckman Optima analytical ultracentrifuge, both equipped with UV-visible scanning optics. Samples were loaded into double-sector cells with quartz windows, and centrifuged using an An-60 Ti or An-50 Ti rotor at 50,000 rpm and at 20 °C. Radial absorbance data was collected in continuous mode at 230, 250 or 280 nm. Sedimentation data were fitted to a continuous sedimentation coefficient c(*s*) model, with floating frictional ratios using *SEDFIT*^99^. Buffer density, viscosity and the partial specific volume of the protein samples were calculated using *SEDNTERP*^100^. For complexes, the partial specific volume used was 0.73 mL/g.

SE-AUC experiments were conducted using a Beckman Coulter XL-I analytical ultracentrifuge, equipped with UV-visible scanning optics. 160 μL of sample was loaded into double-sector cells and centrifuged using an An-60 TI rotor. To calculate M* for each component of the IL-11 signalling complex, proteins were diluted such that A_280_ ∼0.35, and then centrifuged sequentially at 10,500 rpm, 17,000 rpm and 28,000 rpm until equilibrium was reached. For gp130_EC_, the three speeds were 8000 rpm, 10,000 rpm and 16,000 rpm. For the IL-11_Δ10_/gp130_D1-D3_/IL-11Rα_D1-D3_ complex the complex was diluted such at A_250_ ∼0.35 and then centrifuged sequentially at 5,300 rpm, 6,300 rpm and 9,000 rpm. For the IL-11_Δ10_/gp130_EC_/IL-11Rα_D1-D3_ complex, the three speeds used were 4000 rpm, 5000 rpm and 8000 rpm. For all samples, *M*\* was determined using a single-species analysis model in *SEDPHAT*^*101*^. For the complexes, the sample was size exclusion purified prior to analysis, and sample purity was confirmed with SV-AUC prior to the SE-AUC experiment.

### Multi-angle light scattering

SEC-MALS data were collected using a Shimadzu LC-20AD HPLC, coupled to a Shimadzu SPD-20A UV detector, Wyatt Dawn MALS detector and Wyatt Optilab refractive index detector. Data were collected following in-line fractionation with a Zenix-C SEC-300 4.6 × 300 mm SEC column (Sepax Technologies), pre-equilibrated in 20 mM Tris, 150 mM sodium chloride pH 8.5, running at a flow rate of 0.35 mL/min. 10 µL of sample was applied to the column at a concentration of approximately 2 mg/mL. MALS data were analysed using ASTRA v.7.3.2.19 (Wyatt). The MALS detector response was normalised using monomeric bovine serum albumin (BSA) (Pierce, cat no. 23209). Protein concentration was determined using differential refractive index, using a dn/dc of 0.184.

### Small-angle X-ray scattering

SAXS experiments were conducted at the Australian Synchrotron SAXS/WAXS beamline^102-104^. The X-ray beam energy was 11,500 eV (λ = 1.078 Å), the sample-to-detector distance is noted in Supplementary Table 3. Data were collected following fractionation with an in-line size-exclusion chromatography column (Superdex 200 5/150 Increase, GE Healthcare,) pre-equilibrated in TBS pH 8.5, 0.2 % sodium azide. Data were reduced and analysed using Scatterbrain, CHROMIXS^105^ and ATSAS^105,106^, data were analysed using CRYSOL and DAMMIF/DAMAVER/DAMMIN^107-109^. A summary is given in Supplementary Table 3.

Models were prepared of the IL-11_Δ10_/IL-11Rα_D1-D3_/gp130_D1-D3_ complex, the IL-11_Δ10_/IL-11Rα_D1-D3_/gp130_D2-D3_ complex, the IL-11_Δ10_/IL-11Rα_D1-D3_/gp130_EC_ complex, and the IL-11_Δ10/Mutein_/IL-11Rα_D1-D3_/gp130_D1-D3_ complex based on the IL-11_Δ10_/IL-11Rα_D1-D3_/gp130_EC_ complex (see Supplementary Table 3 for the regions of the complex used) for fitting to scattering data in *CRYSOL*. To fit the IL-11_Δ10_/IL-11Rα_D1-D3_/gp130_EC_ complex data, the model of the complex with the D5-D6 domains included was used. For the crystal structure of the IL-11 signalling complex, one hexamer from the crystal structure was used. For IL-11_Δ10/Mutein_ and IL-11_Δ10/W147A_, the unmodified crystal structure coordinates were used.

### Isothermal titration calorimetry

Protein samples were buffer exchanged into TBS pH 8.5 using size exclusion prior to analysis by ITC. ITC data was collected using a MicroCal iTC 200 (GE Healthcare). Titrations were performed using 15 2.5 μL injections of the cytokine ligand, after an initial injection of 0.8 μL. IL-11Rα_D1-D3_ was present at a concentration of approximately 10 μM and the concentration of IL-11 or the relevant IL-11 mutant in the syringe was 10-fold greater than the concentration of IL-11Rα. Following the formation of the cytokine/receptor complex, gp130_D1-D3_ or gp130_D2-D3_ was loaded in the syringe at a concentration approximately 10-fold greater than the concentration of IL-11Rα in the cell, a subsequent titration of gp130 against the cytokine/IL-11Rα_D1-D3_ complex was then performed at 288 K. Titration data were integrated using *NITPIC*^110,111^, and analysed in *SEDPHAT* using a 1:1 interaction model^112^. Each titration was conducted in triplicate, values stated are the mean ± standard error of the mean. Significance of differences were calculated using a one-tailed paired t-test in Microsoft Excel.

### Cell stimulations and phospho-Stat3 flow cytometry assay

Human gp130/IL-11Rα Ba/F3 cells were cultured in RPMI-1640 Medium containing 10% v/v heat inactivated FBS, 1% v/v Penicillin-Streptomycin and 10 ng/mL recombinant IL-11_Δ10_ at 37°C and 5% CO_2_. Cancer cell lines were cultured in RPMI-1640 Medium containing 10% v/v heat inactivated FBS, 1% v/v Penicillin-Streptomycin, except for MDA-MD-231 which were cultured in DMEM. Each cell line was routinely tested for mycoplasma and confirmed to be negative prior to experiments.

Cells were seeded at 50,000 cells per well of a 96-well plate overnight and serum starved for 2 hours prior to stimulation. To determine EC_50_s, the cells were stimulated with the indicated concentration of IL-11_Δ10_, IL-11_Δ10/Mutein_, IL-11_Δ10/W147A_, or IL-11_Δ10/PAIDY_ prepared in Fixable Viability Dye for 15 minutes at 37°C. To determine IC_50_s, the cells were incubated with the indicated concentration of 11_Δ10/Mutein_, IL-11_Δ10/W147A_, or IL-11_Δ10/PAIDY_ for 15 minutes at 37 °C followed by stimulation with 20 ng/mL IL-11_Δ10_ prepared in Fixable Viability Dye and incubated for a further 15 minutes at 37°C.

Cells were harvested by centrifugation at 1600 rpm for 3 minutes and fixed in Cytofix Fixation Buffer (BD) for 10 minutes at 37 °C. Cells were then centrifuged and washed with Stain Buffer and permeabilised with Phosphoflow Perm buffer III for 30 minutes at 40°C. Cells were then centrifuged, washed with Stain Buffer and stained for p-STAT3 (clone: 4/P-STAT3, BD). Data was acquired on a BD Fortessa instrument and analysed using FloJo software. Experiments were conducted in triplicate and EC_50_/IC_50_ values are presented as mean ± standard error of the mean.

### Molecular dynamics

All MD simulations were performed using NAMD 2.1.3b1^113^ and the CHARMM22 forcefield^113,114^ at 310 K in a water box with periodic boundary conditions. Simulations were analysed in VMD 1.9.3^115^. A model of each of IL-11_Δ10_ and IL-11_Δ10/Mutein_ was built based on the crystal structures, for residues with multiple orientations, only one was selected. The structures were solvated (box size 53.6 × 53.1 × 74.9 Å), and ions added to approximate final concentration of 0.15 M NaCl. The MD simulations was performed using a 10 ps minimisation time, followed by 1050 ns MD.

Visualisation and analysis was performed in VMD 1.9.3^115^. A script was used in VMD to measure the distance and calculate the distance distribution between T56/S53 Oγ and H86 Nε throughout the simulation. A script was used to calculate the hydrogen bond potential energy and potential energy distribution of this interaction using a simple electrostatic model, based on the model used in the secondary structure analysis program *DSSP*^116^. Per-residue Cα RMSD and order parameter (*S*^2^) values were calculated using scripts in VMD.

### Differential scanning fluorometry

Protein samples were analysed by DSF at a concentration of 0.1 mg/mL in TBS pH 8 + 0.02% sodium azide, with 2.5 × SYPRO Orange dye (Sigma Aldrich). 20 μL of the sample was loaded into 96-well qPCR plate (Applied Biosystems), and four technical replicates of each sample were analysed. The plates were sealed, and samples heated in an Applied Biosystems StepOne Plus qPCR instrument, from 20 °C to 95 °C, with a 1% gradient. Unfolding data were analysed using a custom script in MATLAB r2019a. The temperature of hydrophobic exposure (*T*_h_), was defined as the minimum point of the first derivative curve, and used to compare the thermal stability of different proteins^68^. All experiments were repeated three times, values are given as mean ± standard error. Significance of differences were calculated using a two-tailed paired t-test in Microsoft Excel.

### Surface plasmon resonance

SPR experiments were conducted using a Biacore T200, at 25 °C, in TBS pH 8.5 + 0.05% Tween, running at 30 μL/min. Biotinylated IL-11_Δ10_-avi and IL-11_Δ10/Mutein_-avi were loaded onto separate channels on a SAHC 1500M streptavidin chip (Xantec). Both proteins were immobilised until *R*_max_ was approximately 70. The chip was washed extensively with 1 M NaCl until the baseline was stable. A nine-point, two-fold dilution series of IL-11Rα_D1-D3_ was prepared, starting at a concentration of 500 nM. Each IL-11Rα dilution was injected over both flow cells in triplicate and reference-subtracted data was generated by subtracting the response from channels in which no protein was loaded from channels containing protein. Data were fitted to a 1:1 kinetic model and the Biacore analysis software was used to determine association (*k*_a_) and dissociation (*k*_d_) rates, and the dissociation constant, *K*_D_. Each dilution series was prepared and analysed in duplicate; values are given as mean ± standard error.

## References

1 Anderson, K. C. et al. Interleukin-11 promotes accessory cell-dependent B-cell differentiation in humans. Blood 80, 2797–2804 (1992).

2 Curti, A. et al. Interleukin-11 induces Th2 polarization of human CD4(+) T cells. Blood 97, 2758–2763, doi:10.1182/blood.v97.9.2758 (2001).

3 Zhang, X. et al. IL-11 Induces Th17 Cell Responses in Patients with Early Relapsing-Remitting Multiple Sclerosis. J Immunol 194, 5139–5149, doi:10.4049/jimmunol.1401680 (2015).

4 Elshabrawy, H. A. et al. IL-11 facilitates a novel connection between RA joint fibroblasts and endothelial cells. Angiogenesis 21, 215–228, doi:10.1007/s10456-017-9589-y (2018).

5 Huynh, J. et al. Host IL11 Signaling Suppresses CD4(+) T cell-Mediated Antitumor Responses to Colon Cancer in Mice. Cancer Immunol Res 9, 735–747, doi:10.1158/2326-6066.CIR-19-1023 (2021).

6 Nayar, S. et al. A myeloid-stromal niche and gp130 rescue in NOD2-driven Crohn’s disease. Nature, doi:10.1038/s41586-021-03484-5 (2021).

7 Minshall, E. et al. IL-11 expression is increased in severe asthma: association with epithelial cells and eosinophils. J Allergy Clin Immunol 105, 232–238, doi:10.1016/s0091-6749(00)90070-8 (2000).

8 Calon, A. et al. Dependency of colorectal cancer on a TGF-beta-driven program in stromal cells for metastasis initiation. Cancer Cell 22, 571–584, doi:10.1016/j.ccr.2012.08.013 (2012).

9 Ng, B. et al. Interleukin-11 is a therapeutic target in idiopathic pulmonary fibrosis. Sci Transl Med 11, doi:10.1126/scitranslmed.aaw1237 (2019).

10 Schafer, S. et al. IL-11 is a crucial determinant of cardiovascular fibrosis. Nature 552, 110–115, doi:10.1038/nature24676 (2017).

11 Widjaja, A. A. et al. Inhibiting Interleukin 11 Signaling Reduces Hepatocyte Death and Liver Fibrosis, Inflammation, and Steatosis in Mouse Models of Nonalcoholic Steatohepatitis. Gastroenterology 157, 777–792 e714, doi:10.1053/j.gastro.2019.05.002 (2019).

12 Paul, S. R. et al. Molecular cloning of a cDNA encoding interleukin 11, a stromal cell-derived lymphopoietic and hematopoietic cytokine. Proceedings of the National Academy of Sciences 87, 7512–7516, doi:10.1073/pnas.87.19.7512 (1990).

13 Metcalfe, R., Putoczki, T. & Griffin, M. Structural Understanding of Interleukin 6 Family Cytokine Signaling and Targeted Therapies: Focus on Interleukin 11. Frontiers in Immunology 11, 1–25, doi:10.3389/fimmu.2020.01424 (2020).

14 Wu, S. et al. Multicenter, randomized study of genetically modified recombinant human interleukin-11 to prevent chemotherapy-induced thrombocytopenia in cancer patients receiving chemotherapy. Support Care Cancer 20, 1875–1884, doi:10.1007/s00520-011-1290-x (2012).

15 Okamoto, H. et al. The synovial expression and serum levels of interleukin-6, interleukin-11, leukemia inhibitory factor, and oncostatin M in rheumatoid arthritis. Arthritis Rheum 40, 1096–1105, doi:10.1002/art.1780400614 (1997).

16 Adami, E. et al. IL11 is elevated in systemic sclerosis and IL11-dependent ERK signaling underlies TGFbeta-mediated activation of dermal fibroblasts. Rheumatology (Oxford), doi:10.1093/rheumatology/keab168 (2021).

17 Strikoudis, A. et al. Modeling of Fibrotic Lung Disease Using 3D Organoids Derived from Human Pluripotent Stem Cells. Cell Reports, 3709–3723, doi:10.1016/j.celrep.2019.05.077 (2019).

18 Corden, B., Adami, E., Sweeney, M., Schafer, S. & Cook, S. A. IL-11 in cardiac and renal fibrosis: Late to the party but a central player. Br J Pharmacol 177, 1695–1708, doi:10.1111/bph.15013 (2020).

19 Nayar, S. et al. A myeloid-stromal niche and gp130 rescue in NOD2-driven Crohn’s disease. Nature 593, 275–281, doi:10.1038/s41586-021-03484-5 (2021).

20 Corden, B. et al. Therapeutic Targeting of Interleukin-11 Signalling Reduces Pressure Overload-Induced Cardiac Fibrosis in Mice. J Cardiovasc Transl Res 14, 222–228, doi:10.1007/s12265-020-10054-z (2021).

21 Widjaja, A. A. et al. A Neutralizing IL-11 Antibody Improves Renal Function and Increases Lifespan in a Mouse Model of Alport Syndrome. J Am Soc Nephrol 33, 718–730, doi:10.1681/ASN.2021040577 (2022).

22 van Duijneveldt, G., Griffin, M. D. W. & Putoczki, T. L. Emerging roles for the IL-6 family of cytokines in pancreatic cancer. Clinical Science 134, 2091–2115, doi:10.1042/CS20191211 (2020).

23 Liang, M. et al. IL-11 is essential in promoting osteolysis in breast cancer bone metastasis via RANKL-independent activation of osteoclastogenesis. Cell Death Dis 10, 353, doi:10.1038/s41419-019-1594-1 (2019).

24 Ernst, M. et al. STAT3 and STAT1 mediate IL-11-dependent and inflammation-associated gastric tumorigenesis in gp130 receptor mutant mice. J Clin Invest 118, 1727–1738, doi:10.1172/JCI34944 (2008).

25 Putoczki, T. L. et al. Interleukin-11 is the dominant IL-6 family cytokine during gastrointestinal tumorigenesis and can be targeted therapeutically. Cancer Cell 24, 257–271, doi:10.1016/j.ccr.2013.06.017 (2013).

26 Lay, V., Yap, J., Sonderegger, S. & Dimitriadis, E. Interleukin 11 regulates endometrial cancer cell adhesion and migration via STAT3. Int J Oncol 41, 759–764, doi:10.3892/ijo.2012.1486 (2012).

27 Barton, V., Hall, M., Hudson, K. R. & Heath, J. K. Interleukin-11 signals through the formation of a hexameric receptor complex. Journal of Biological Chemistry 275, 36197–36203, doi:10.1074/jbc.M004648200 (2000).

28 Morris, R., Kershaw, N. J. & Babon, J. J. The molecular details of cytokine signaling via the JAK/STAT pathway. Protein Science 27, 1984–2009, doi:10.1002/pro.3519 (2018).

29 Mahboubi, K., Biedermann, B. C., Carroll, J. M. & Pober, J. S. IL-11 activates human endothelial cells to resist immune-mediated injury. J Immunol 164, 3837–3846, doi:10.4049/jimmunol.164.7.3837 (2000).

30 Thiem, S. et al. mTORC1 inhibition restricts inflammation-associated gastrointestinal tumorigenesis in mice. J Clin Invest 123, 767–781, doi:10.1172/JCI65086 (2013).

31 Derouet, D. et al. Neuropoietin, a new IL-6-related cytokine signaling through the ciliary neurotrophic factor receptor. Proc Natl Acad Sci U S A 101, 4827–4832, doi:10.1073/pnas.0306178101 (2004).

32 Murakami, M., Kamimura, D. & Hirano, T. New IL-6 (gp130) family cytokine members, CLC/NNT1/BSF3 and IL-27. Growth Factors 22, 75–77, doi:10.1080/08977190410001715181 (2004).

33 Rose-John, S. Interleukin-6 Family Cytokines. Cold Spring Harb Perspect Biol 10, doi:10.1101/cshperspect.a028415 (2018).

34 Tait Wojno, E. D., Hunter, C. A. & Stumhofer, J. S. The Immunobiology of the Interleukin-12 Family: Room for Discovery. Immunity 50, 851–870, doi:10.1016/j.immuni.2019.03.011 (2019).

35 Chow, D.-C., He, X., Snow, a. L., Rose-John, S. & Garcia, K. Structure of an extracellular gp130 cytokine receptor signaling complex. Science 291, 2150–2155, doi:10.1126/science.1058308 (2001).

36 Boulanger, M. J., Chow, D.-c., Brevnova, E. E. & Garcia, K. C. Hexameric structure and assembly of the interleukin-6/IL-6 alpha-receptor/gp130 complex. Science 300, 2101–2104, doi:10.1126/science.1083901 (2003).

37 Huyton, T. et al. An unusual cytokine:Ig-domain interaction revealed in the crystal structure of leukemia inhibitory factor (LIF) in complex with the LIF receptor. Proceedings of the National Academy of Sciences 104, 12737–12742, doi:10.1073/pnas.0705577104 (2007).

38 Boulanger, M. J., Bankovich, A. J., Kortemme, T., Baker, D. & Garcia, K. C. Convergent mechanisms for recognition of divergent cytokines by the shared signaling receptor gp130. Molecular Cell 12, 577–589, doi:10.1016/S1097-2765(03)00365-4 (2003).

39 Wong, P. K., Campbell, I. K., Robb, L. & Wicks, I. P. Endogenous IL-11 is pro-inflammatory in acute methylated bovine serum albumin/interleukin-1-induced (mBSA/IL-1)arthritis. Cytokine 29, 72–76, doi:10.1016/j.cyto.2004.09.011 (2005).

40 Schumacher, D. et al. A neutralizing IL-11 antibody reduces vessel hyperplasia in a mouse carotid artery wire injury model. Sci Rep 11, 20674, doi:10.1038/s41598-021-99880-y (2021).

41 Chun, G. L. et al. Endogenous IL-11 signaling is essential in Th2-and IL-13-induced inflammation and mucus production. American Journal of Respiratory Cell and Molecular Biology 39, 739–746, doi:10.1165/rcmb.2008-0053OC (2008).

42 Underhill-Day, N. et al. Functional characterization of W147A: A high-affinity interleukin-11 antagonist. Endocrinology 144, 3406–3414, doi:10.1210/en.2002-0144 (2003).

43 Wu, P. et al. IL-11 Is Elevated and Drives the Profibrotic Phenotype Transition of Orbital Fibroblasts in Thyroid-Associated Ophthalmopathy. Front Endocrinol (Lausanne) 13, 846106, doi:10.3389/fendo.2022.846106 (2022).

44 Winship, A. L., Van Sinderen, M., Donoghue, J., Rainczuk, K. & Dimitriadis, E. Targeting interleukin-11 receptor-α impairs human endometrial cancer cell proliferation and invasion in vitro and reduces tumor growth and metastasis in vivo. Molecular Cancer Therapeutics 15, 720–730, doi:10.1158/1535-7163.MCT-15-0677 (2016).

45 Ng, B. et al. IL11 Activates Pancreatic Stellate Cells and Causes Pancreatic Inflammation, Fibrosis and Atrophy in a Mouse Model of Pancreatitis. Int J Mol Sci 23, doi:10.3390/ijms23073549 (2022).

46 Lim, W. W. et al. Inhibition of IL11 Signaling Reduces Aortic Pathology in Murine Marfan Syndrome. Circ Res 130, 728–740, doi:10.1161/CIRCRESAHA.121.320381 (2022).

47 Metcalfe, R. D. et al. The structure of the extracellular domains of human interleukin 11alpha receptor reveals mechanisms of cytokine engagement. J Biol Chem 295, 8285–8301, doi:10.1074/jbc.RA119.012351 (2020).

48 Matadeen, R., Hon, W. C., Heath, J. K., Jones, E. Y. & Fuller, S. The Dynamics of Signal Triggering in a gp130-Receptor Complex. Structure 15, 441–448, doi:10.1016/j.str.2007.02.006 (2007).

49 Skiniotis, G., Boulanger, M. J., Garcia, K. C. & Walz, T. Signaling conformations of the tall cytokine receptor gp130 when in complex with IL-6 and IL-6 receptor. Nature Structural and Molecular Biology 12, 545–551, doi:10.1038/nsmb941 (2005).

50 Lupardus, P. J. et al. Structural snapshots of full-length Jak1, a transmembrane gp130/IL-6/IL-6Rα cytokine receptor complex, and the receptor-Jak1 holocomplex. Structure 19, 45–55, doi:10.1016/j.str.2010.10.010 (2011).

51 Zhou, Y. et al. Structural insights into the assembly of gp130 family cytokine signaling complexes. bioRxiv, doi:10.1101/2022.06.30.496838 (2022).

52 Barton, V., Hudson, K. R. & Heath, J. K. Identification of three distinct receptor binding sites of murine interleukin-11. Journal of Biological Chemistry 274, 5755–5761, doi:10.1074/jbc.274.9.5755 (1999).

53 Tacken, I. et al. Definition of receptor binding sites on human interleukin-11 by molecular modeling-guided mutagenesis. European Journal of Biochemistry 265, 645–655, doi:10.1046/j.1432-1327.1999.00755.x (1999).

54 Czupryn, M. et al. Alanine-scanning mutagenesis of human interleukin-11: identification of regions important for biological activity. Annals of the New York Academy of Sciences 762, 152–164 (1995).

55 Kurth, I. et al. Activation of the signal transducer glycoprotein 130 by both IL-6 and IL-11 requires two distinct binding epitopes. Journal of immunology 162, 1480–1487 (1999).

56 Li, H. & Nicholas, J. Identification of Amino Acid Residues of gp130 Signal Transducer and gp80 α Receptor Subunit That Are Involved in Ligand Binding and Signaling by Human Herpesvirus 8-Encoded Interleukin-6. Journal of Virology 76, 5627–5636, doi:10.1128/jvi.76.11.5627-5636.2002 (2002).

57 Xu, Y. et al. Crystal structure of the entire ectodomain of gp130: Insights into the molecular assembly of the tall cytokine receptor complexes. Journal of Biological Chemistry 285, 21214–21218, doi:10.1074/jbc.C110.129502 (2010).

58 Skiniotis, G., Lupardus, P. J., Martick, M., Walz, T. & Garcia, K. C. Structural Organization of a Full-Length gp130/LIF-R Cytokine Receptor Transmembrane Complex. Molecular Cell 31, 737–748, doi:10.1016/j.molcel.2008.08.011 (2008).

59 Timmermann, A., Küster, A., Kurth, I., Heinrich, P. C. & Müller-Newen, G. A functional role of the membrane-proximal extracellular domains of the signal transducer gp130 in heterodimerization with the leukemia inhibitory factor receptor. European Journal of Biochemistry 269, 2716–2726, doi:10.1046/j.1432-1033.2002.02941.x (2002).

60 Kurth, I. et al. Importance of the Membrane-Proximal Extracellular Domains for Activation of the Signal Transducer Glycoprotein 130. The Journal of Immunology 164, 273–282, doi:10.4049/jimmunol.164.1.273 (2000).

61 Punjani, A. & Fleet, D. J. 3D variability analysis: Resolving continuous flexibility and discrete heterogeneity from single particle cryo-EM. Journal of Structural Biology 213, 107702, doi:10.1016/j.jsb.2021.107702 (2021).

62 Punjani, A., Rubinstein, J. L., Fleet, D. J. & Brubaker, M. A. CryoSPARC: Algorithms for rapid unsupervised cryo-EM structure determination. Nature Methods 14, 290–296, doi:10.1038/nmeth.4169 (2017).

63 Chen, Y. H. et al. Functional and structural analysis of cytokine-selective IL6ST defects that cause recessive hyper-IgE syndrome. J Allergy Clin Immunol 148, 585–598, doi:10.1016/j.jaci.2021.02.044 (2021).

64 Nandurkar, H. H. et al. The human IL-11 receptor requires gp130 for signalling: demonstration by molecular cloning of the receptor. Oncogene 12, 585–593 (1996).

65 Shepelkova, G., Evstifeev, V., Majorov, K., Bocharova, I. & Apt, A. Therapeutic Effect of Recombinant Mutated Interleukin 11 in the Mouse Model of Tuberculosis. J Infect Dis 214, 496–501, doi:10.1093/infdis/jiw176 (2016).

66 Wang, H., Wang, D. H., Yang, X., Sun, Y. & Yang, C. S. Colitis-induced IL11 promotes colon carcinogenesis. Carcinogenesis 42, 557–569, doi:10.1093/carcin/bgaa122 (2021).

67 Ashraf, S. S., Benson, R. E., Payne, E. S., Halbleib, C. M. & Grøn, H. A novel multi-affinity tag system to produce high levels of soluble and biotinylated proteins in Escherichia coli. Protein Expression and Purification 33, 238–245, doi:10.1016/j.pep.2003.10.016 (2004).

68 Seabrook, S. a. & Newman, J. High-throughput thermal scanning for protein stability: Making a good technique more robust. ACS Combinatorial Science 15, 387–392, doi:10.1021/co400013v (2013).

69 Widjaja, A. A. et al. Molecular Dissection of Pro-Fibrotic IL11 Signaling in Cardiac and Pulmonary Fibroblasts. Front Mol Biosci 8, 740650, doi:10.3389/fmolb.2021.740650 (2021).

70 Chen, Y. H. et al. Absence of GP130 cytokine receptor signaling causes extended Stüve-Wiedemann syndrome. Journal of Experimental Medicine 217, doi:10.1084/jem.20191306 (2020).

71 Béziat, V. et al. Dominant-negative mutations in human IL6ST underlie hyper-IgE syndrome. Journal of Experimental Medicine 217, doi:10.1084/jem.20191804 (2020).

72 Shahin, T. et al. Selective loss of function variants in IL6ST cause hyper-IgE syndrome with distinct impairments of T-cell phenotype and function. Haematologica 104, 609–621, doi:10.3324/haematol.2018.194233 (2019).

73 Schwerd, T. et al. A biallelic mutation in IL6ST encoding the GP130 coreceptor causes immunodeficiency and craniosynostosis. Journal of Experimental Medicine 214, 2547–2562, doi:10.1084/jem.20161810 (2017).

74 Monies, D. et al. Lessons Learned from Large-Scale, First-Tier Clinical Exome Sequencing in a Highly Consanguineous Population. American Journal of Human Genetics 104, 1182–1201, doi:10.1016/j.ajhg.2019.04.011 (2019).

75 Schwerd, T. et al. A variant in IL6ST with a selective IL-11 signaling defect in human and mouse. Bone Research 8, doi:10.1038/s41413-020-0098-z (2020).

76 Nieminen, P. et al. Inactivation of IL11 signaling causes craniosynostosis, delayed tooth eruption, and supernumerary teeth. American Journal of Human Genetics 89, 67–81, doi:10.1016/j.ajhg.2011.05.024 (2011).

77 Neveling, K. et al. Mutations in the interleukin receptor IL11RA cause autosomal recessive Crouzon-like craniosynostosis. Molecular Genetics and Genomic Medicine 1, 223–237, doi:10.1002/mgg3.28 (2013).

78 Klein, W. et al. A promotor polymorphism in the Interleukin 11 gene is associated with chronic obstructive pulmonary disease. Electrophoresis 25, 804–808, doi:10.1002/elps.200305773 (2004).

79 Putoczki, T. L., Dobson, R. C. J. & Griffin, M. D. W. The structure of human interleukin-11 reveals receptor-binding site features and structural differences from interleukin-6. Acta Crystallographica Section D-Biological Crystallography 3, 2277–2285, doi:10.1107/S1399004714012267 (2014).

80 Kimanius, D., Forsberg, B. O., Scheres, S. H. & Lindahl, E. Accelerated cryo-EM structure determination with parallelisation using GPUs in RELION-2. Elife 5, doi:10.7554/eLife.18722 (2016).

81 Scheres, S. H. Semi-automated selection of cryo-EM particles in RELION-1.3. J Struct Biol 189, 114–122, doi:10.1016/j.jsb.2014.11.010 (2015).

82 Terwilliger, T. C., Sobolev, O. V., Afonine, P. V. & Adams, P. D. Automated map sharpening by maximization of detail and connectivity. Acta Crystallographica Section D Structural Biology 74, 545–559, doi:10.1107/S2059798318004655 (2018).

83 Kucukelbir, A., Sigworth, F. J. & Tagare, H. D. Quantifying the local resolution of cryo-EM density maps. Nature Methods 11, 63–65, doi:10.1038/nmeth.2727 (2014).

84 Tegunov, D. & Cramer, P. Real-time cryo-electron microscopy data preprocessing with Warp. Nat Methods 16, 1146–1152, doi:10.1038/s41592-019-0580-y (2019).

85 Emsley, P., Lohkamp, B., Scott, W. G. & Cowtan, K. Features and development of Coot. Acta Crystallographica Section D-Biological Crystallography 66, 486–501, doi:Doi 10.1107/S0907444910007493 (2010).

86 Afonine, P. V. et al. Real-space refinement in Phenix for cryo-EM and crystallography. Acta Crystallographica Section D Structural Biology 74, 531–544, doi:10.1101/249607 (2018).

87 Pettersen, E. F. et al. UCSF Chimera--a visualization system for exploratory research and analysis. J Comput Chem 25, 1605–1612, doi:10.1002/jcc.20084 (2004).

88 Barad, B. A. et al. EMRinger: side chain–directed model and map validation for 3D cryo-electron microscopy. Nature Methods 12, 943–946, doi:10.1038/nmeth.3541 (2015).

89 Williams, C. J. et al. MolProbity: More and better reference data for improved all-atom structure validation. Protein Science 27, 293–315, doi:10.1002/pro.3330 (2018).

90 Shindyalov, I. N. & Bourne, P. E. Protein structure alignment by incremental combinatorial extension (CE) of the optimal path. Protein Engineering 11, 739–747 (1998).

91 Luft, J. R. & DeTitta, G. T. A method to produce microseed stock for use in the crystallization of biological macromolecules. Acta Crystallographica Section D: Biological Crystallography 55, 988–993, doi:10.1107/S0907444999002085 (1999).

92 Aragão, D. et al. MX2: a high-flux undulator microfocus beamline serving both the chemical and macromolecular crystallography communities at the Australian Synchrotron. Journal of Synchrotron Radiation 25, 885–891, doi:10.1107/s1600577518003120 (2018).

93 Kabsch, W. Xds. Acta Crystallographica Section D: Biological Crystallography 66, 125–132, doi:10.1107/S0907444909047337 (2010).

94 Evans, P. Scaling and assessment of data quality. Acta Crystallographica Section D: Biological Crystallography 62, 72–82, doi:10.1107/S0907444905036693 (2006).

95 Evans, P. R. & Murshudov, G. N. How good are my data and what is the resolution? Acta Crystallographica Section D: Biological Crystallography 69, 1204–1214, doi:10.1107/S0907444913000061 (2013).

96 Tickle, I. J. et al. STARANISO, <http://staraniso.globalphasing.org/cgi-bin/staraniso.cgi> (2018).

97 McCoy, A. J. et al. Phaser crystallographic software Journal of Applied Crystallography 40, 658–674, doi:10.1107/s0021889807021206 (2007).

98 Afonine, P. V. et al. Towards automated crystallographic structure refinement with phenix.refine. Acta Crystallographica Section D: Biological Crystallography 68, 352–367, doi:10.1107/S0907444912001308 (2012).

99 Schuck, P. Size-distribution analysis of macromolecules by sedimentation velocity ultracentrifugation and lamm equation modeling. Biophysical Journal 78, 1606–1619, doi:10.1016/S0006-3495(00)76713-0 (2000).

100 Laue, T. M., Shah, B. D., Ridgeway, T. M. & Pelletier, S. L. Analytical Ultracentrifugation in Biochemistry and Polymer Science. 90–125 (Cambridge: Royal Society of Chemistry, 1992).

101 Vistica, J. et al. Sedimentation equilibrium analysis of protein interactions with global implicit mass conservation constraints and systematic noise decomposition. Analytical Biochemistry 326, 234–256, doi:10.1016/j.ab.2003.12.014 (2004).

102 Kirby, N. M. et al. A low-background-intensity focusing small-angle X-ray scattering undulator beamline. Journal of Applied Crystallography 46, 1670–1680, doi:10.1107/S002188981302774X (2013).

103 Kirby, N. et al. Improved radiation dose efficiency in solution SAXS using a sheath flow sample environment. Acta Crystallographica Section D Structural Biology 72, 1254–1266, doi:10.1107/S2059798316017174 (2016).

104 Ryan, T. M. et al. An optimized SEC-SAXS system enabling high X-ray dose for rapid SAXS assessment with correlated UV measurements for biomolecular structure analysis:. Journal of Applied Crystallography 51, 97–111, doi:10.1107/S1600576717017101 (2018).

105 Panjkovich, A. & Svergun, D. I. CHROMIXS: automatic and interactive analysis of chromatography-coupled small-angle X-ray scattering data. Bioinformatics 34, 1944–1946 doi:10.1093/bioinformatics/btx846 (2017).

106 Franke, D. et al. ATSAS 2.8: A comprehensive data analysis suite for small-angle scattering from macromolecular solutions. Journal of Applied Crystallography 50, 1212–1225, doi:10.1107/S1600576717007786 (2017).

107 Franke, D. & Svergun, D. I. DAMMIF, a program for rapid ab-initio shape determination in small-angle scattering. Journal of Applied Crystallography 42, 342–346, doi:10.1107/S0021889809000338 (2009).

108 Volkov, V. V. & Svergun, D. I. Uniqueness of ab initio shape determination in small-angle scattering. Journal of Applied Crystallography 36, 860–864, doi:10.1107/S0021889803000268 (2003).

109 Svergun, D. Restoring low resolution structure of biological macromolecules from solution scattering using simulated annealing. Biophysical Journal 76, 2879–2886 (1999).

110 Brautigam, C. A., Zhao, H., Vargas, C., Keller, S. & Schuck, P. Integration and global analysis of isothermal titration calorimetry data for studying macromolecular interactions. Nature Protocols 11, 882–894, doi:10.1038/nprot.2016.044 (2016).

111 Keller, S. et al. High-Precision Isothermal Titration Calorimetry with Automated Peak Shape Analysis. Analytical Chemistry 84, 5066–5073, doi:10.1021/ac3007522 (2012).

112 Zhao, H. & Schuck, P. Combining biophysical methods for the analysis of protein complex stoichiometry and affinity in SEDPHAT. Acta Crystallographica Section D: Biological Crystallography 71, 3–14, doi:10.1107/S1399004714010372 (2015).

113 Phillips, J. C. et al. Scalable molecular dynamics with NAMD. Journal of Computational Chemistry 26, 1781–1802, doi:10.1002/jcc.20289 (2005).

114 Brooks, B. R. et al. CHARMM: The Biomolecular Simulation Program. Journal of Computational Chemistry 20, 1545–1614, doi:10.1002/jcc (2009).

115 Humphrey, W., Dalke, A. & Schulten, K. VMD: Visual Molecular Dynamics. Journal of Molecular Graphics 7855, 33–38, doi:10.1016/0263-7855(96)00018-5 (1996).

116 Kabsch, W. & Sander, C. Dictonary of Protein Secondary Structure: Pattern Recognition of Hydrogen-Bonded and Geometrical Features. Biopolymers 22, 2578–2637 (1983).

