## Supplemental Information for "Structures of the interleukin 11 signalling complex reveal dynamics of gp130 extracellular domains and the inhibitory mechanism of a cytokine variant"

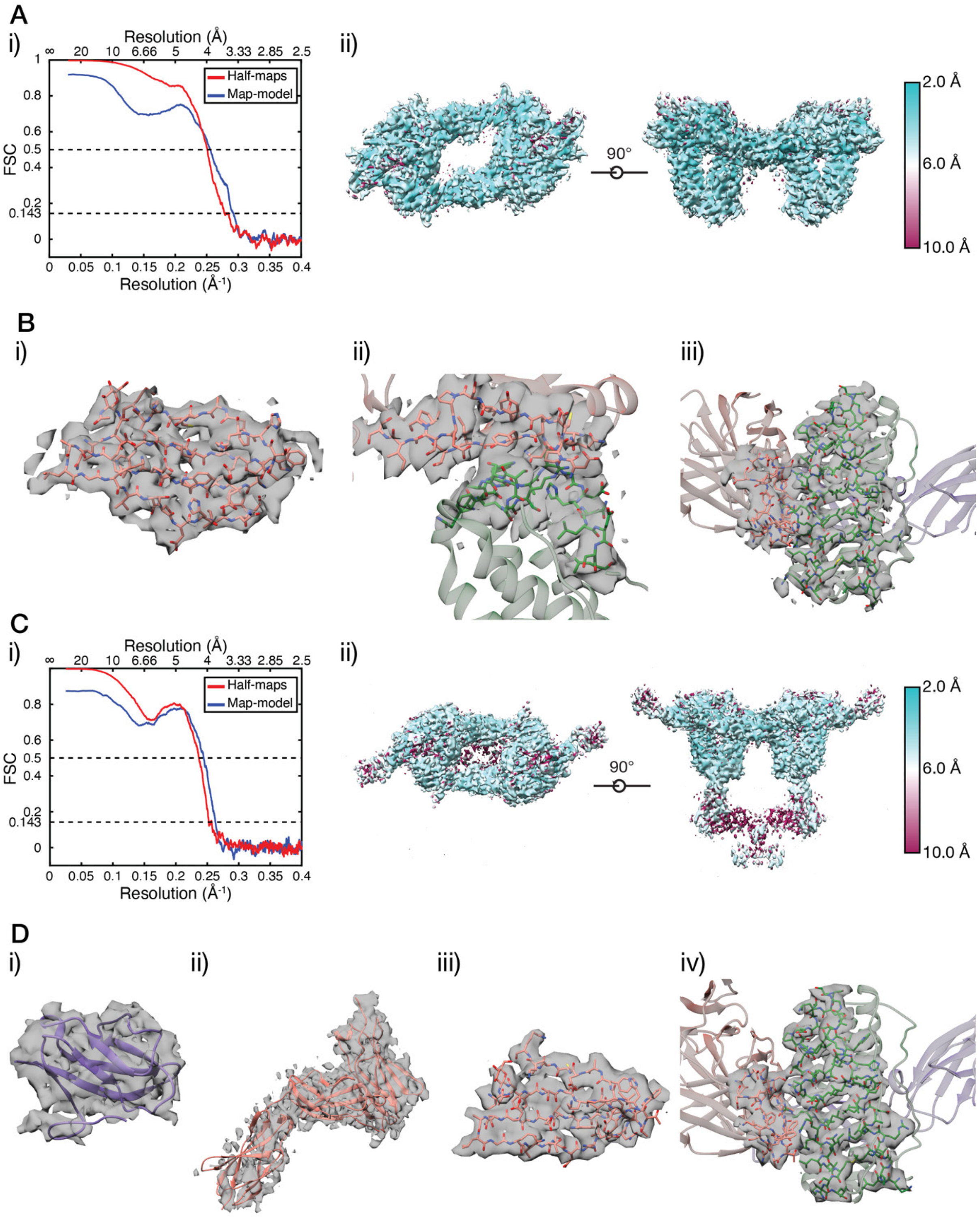

**Supplementary Figure 1:** Resolution estimation and cryo-EM representative density. A) Resolution estimation for the gp130$_{D1\text{-}D3}$ complex, i) map-model and gold-standard half-map Fourier shell correlation (FSC) curves calculated in *Phenix*[1,2], ii) Local resolution maps, calculated using *Resmap*[3]. B) Representative cryo-EM density for the gp130$_{D1\text{-}D3}$ complex, i) one of the β-sheets in gp130 D2, ii) the site-III interface, iii) the site-II interface. C) Resolution estimation for the gp130$_{EC}$ complex, i) map-model and gold-standard half-map Fourier shell correlation (FSC) curves calculated in *Phenix*[1,2], ii) Local resolution maps, calculated using *Resmap*[3]. D) Representative cryo-EM density for the gp130$_{EC}$ complex, i) IL-11Rα D1, ii) gp130 D4-D6, iii) one of the β-sheets in gp130 D2, iv) the site-III interface.

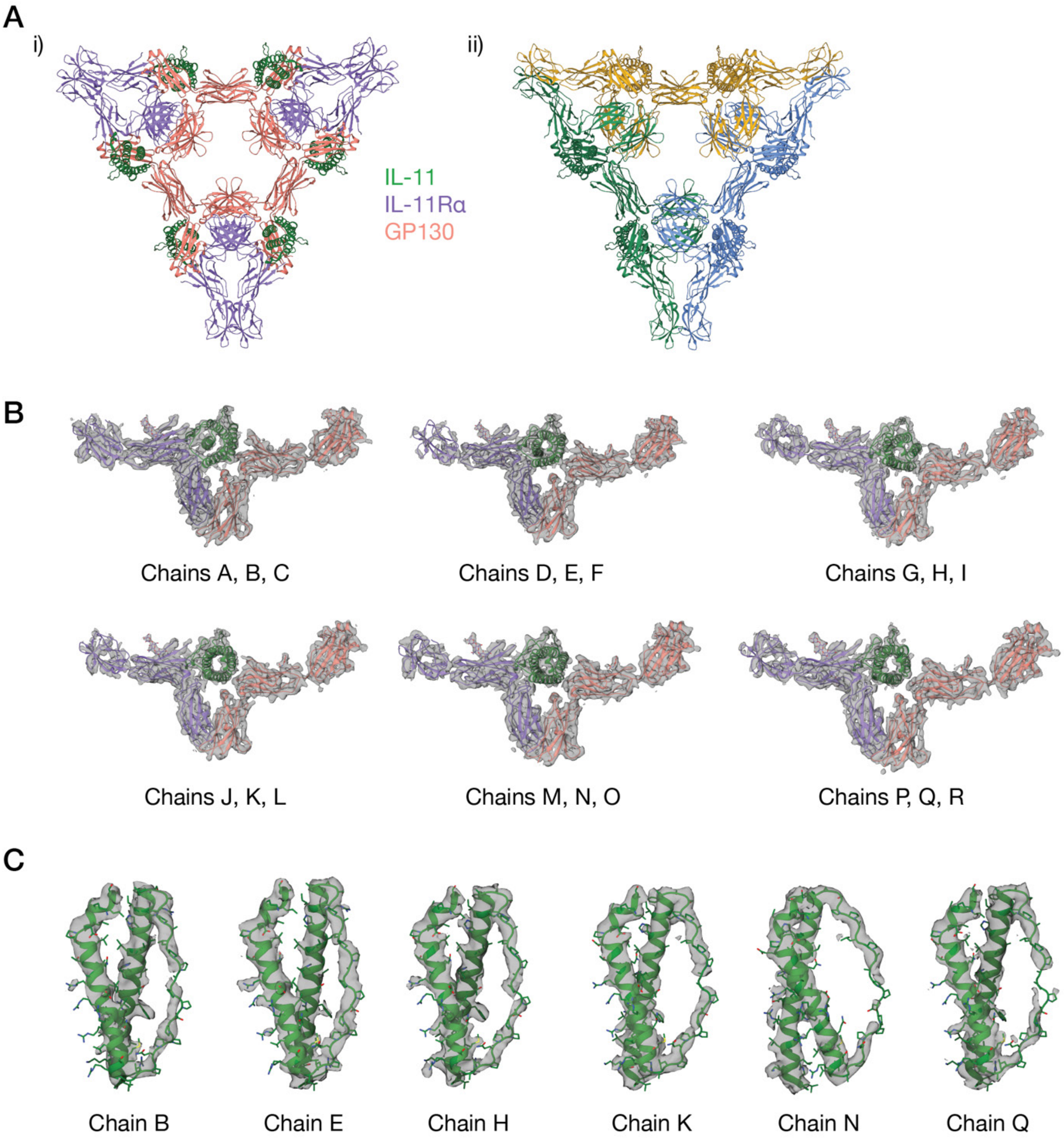

**Supplementary Figure 2:** Asymmetric unit and representative electron density for the IL-11 complex crystal structure. A) The asymmetric unit of the crystal structure of the IL-11 signalling complex, with three hexamers in the asymmetric unit, coloured according to molecule in i), each hexamer coloured differently in ii). B) Representative electron density for each trimer in the crystal structure. C) Representative electron density for the C and D helices in IL-11, from each trimer in the crystal structure. Density contoured at 1 σ, with missing $F_{obs}$ not filled

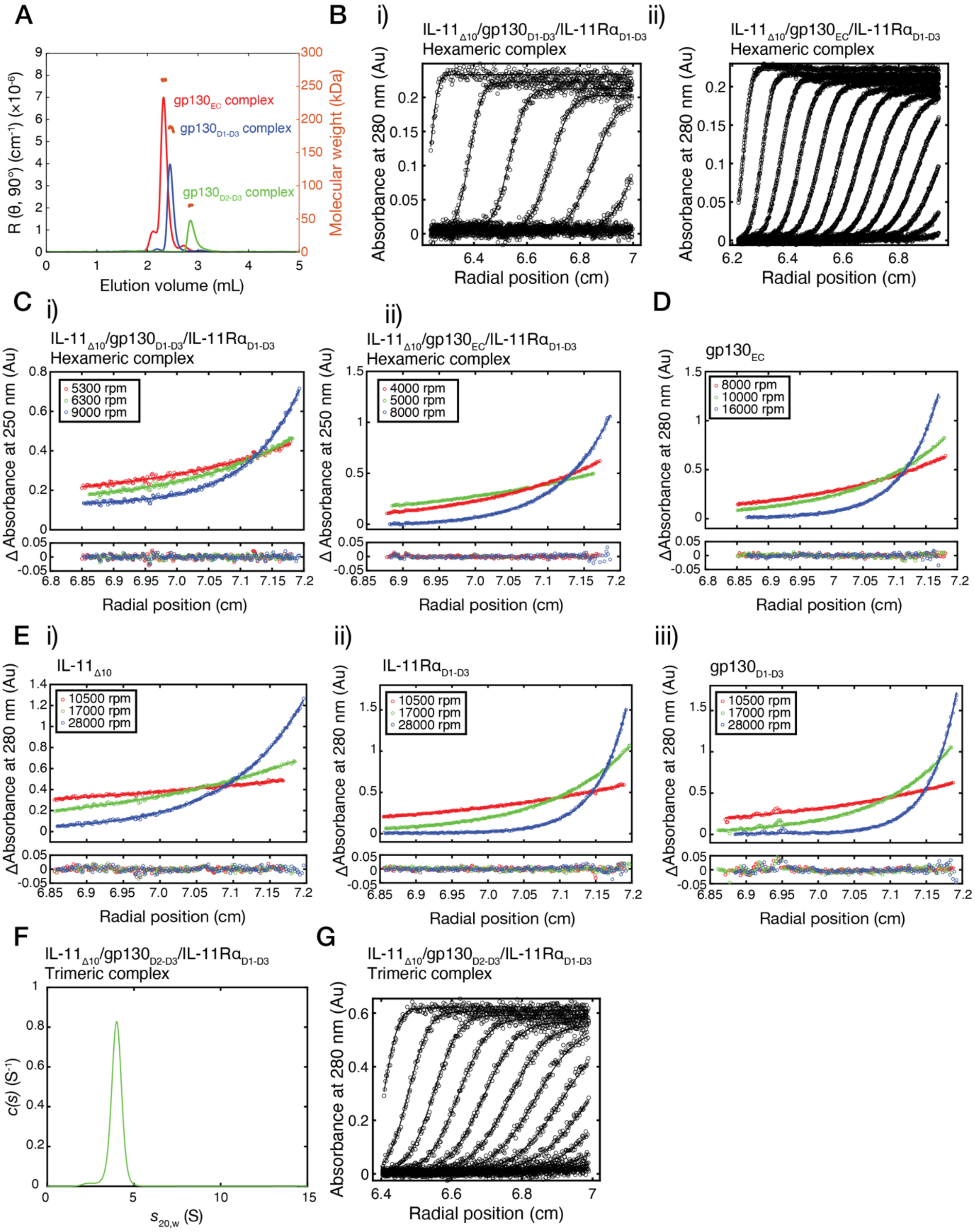

**Supplementary Figure 3:** Raw SV-AUC scans, SE and MALS data. Related to Figure 1C-D. A) Raw SV-AUC scans for the data shown in, i) Figure 1Di, ii) Figure 1Dii, iii) Figure 1Diii. B) SE-AUC data for individual components of the $gp130_{D1\text{-}D3}$ complex, i) $IL\text{-}11_{\Delta10}$, ii) $IL\text{-}11R\alpha_{D1\text{-}D3}$, iii) $gp130_{D1\text{-}D3}$. C) SE-AUC data for $gp130_{EC}$. D) MALS data for the $gp130_{EC}$, $gp130_{D1\text{-}D3}$ and $gp130_{D2\text{-}D3}$ complexes.

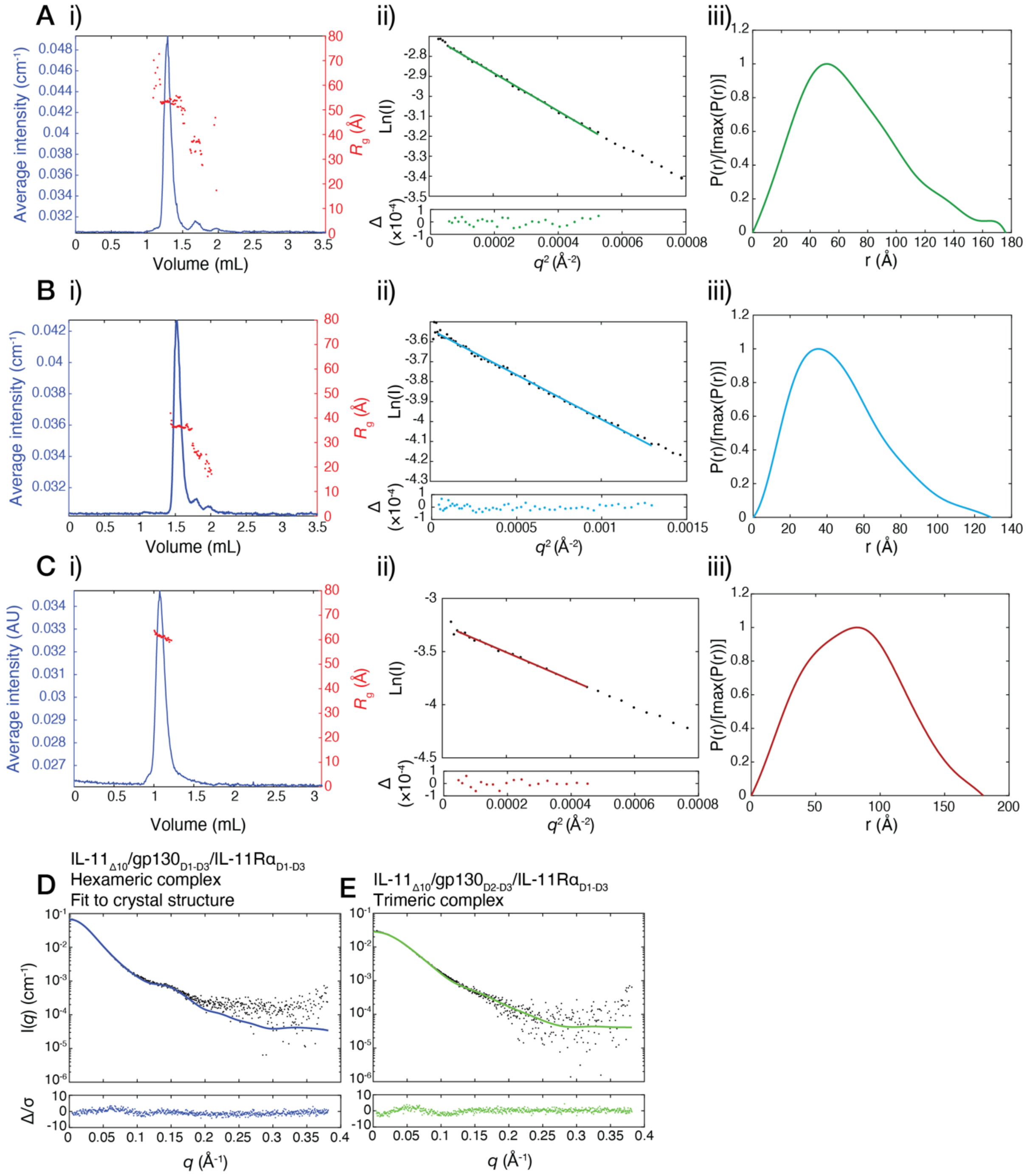

**Supplementary Figure 4:** Supplementary SAXS data, related to Figure 1E. A) Supplemental SAXS data for the gp130$_{D1\text{-}D3}$ complex, i) SEC-SAXS chromatogram, ii) Guinier plot, iii) pairwise distance distribution (P(r)) plot). B) Supplemental SAXS data for the gp130$_{D2\text{-}D3}$ complex, i) SEC-SAXS chromatogram, ii) Guinier plot, iii) pairwise distance distribution (P(r)) plot). C) Supplemental SAXS data for the gp130$_{EC}$ complex, i) SEC-SAXS chromatogram, ii) Guinier plot, iii) pairwise distance distribution (P(r)) plot). D) Fit of one hexamer from the IL-11 signalling complex crystal structure, to the scattering data from the gp130$_{D1\text{-}D3}$ complex (see Supplementary Table 3 for fitting statistics).

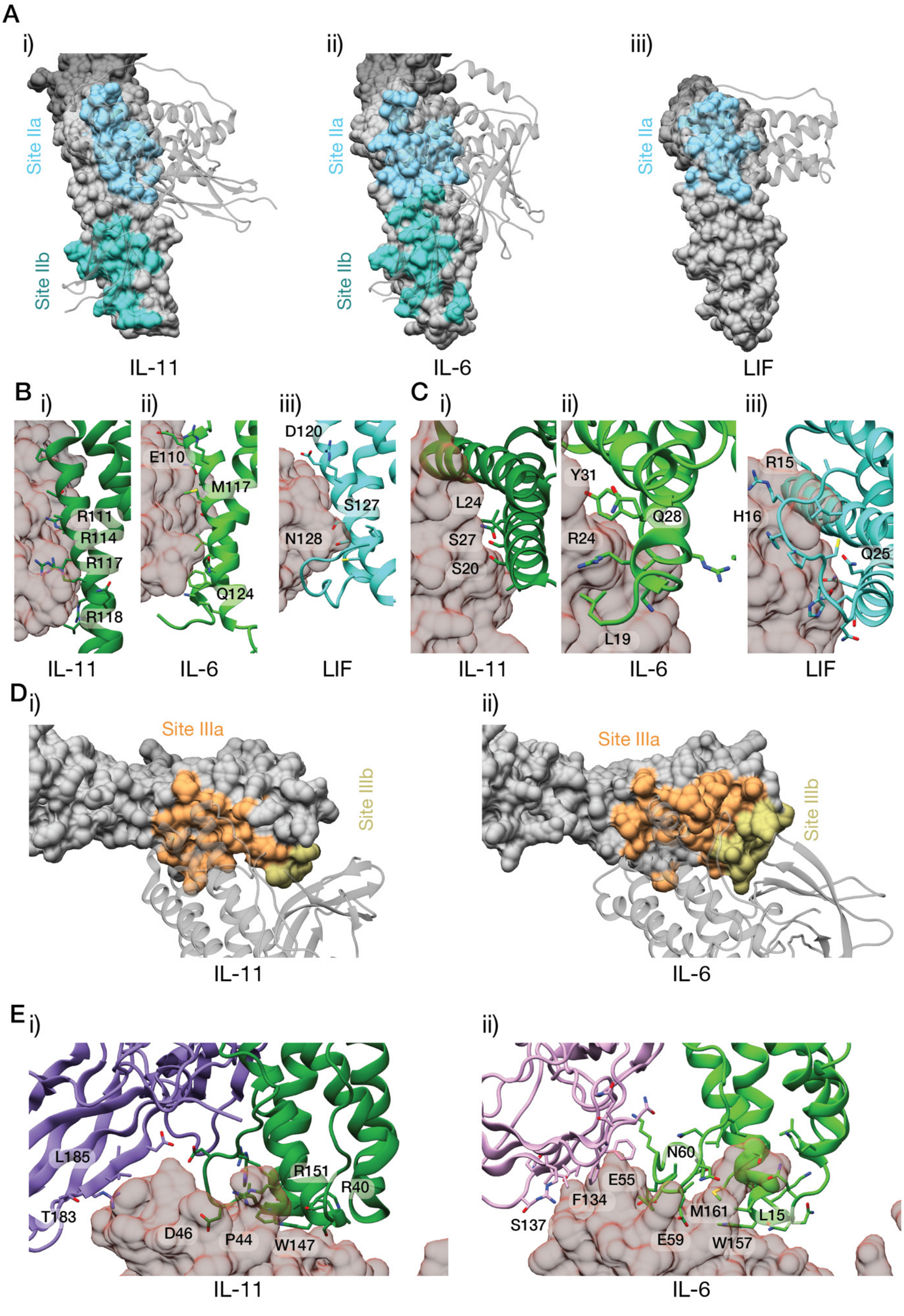

A
i)
ii)
iii)
Site IIa
Site IIb
IL-11
IL-6
LIF
B
C
D120
E110
M117
S127
N128
R111
R114
R117
R118
Q124
L24
S27
S20
Y31
Q28
R24
L19
R15
H16
Q25
D
Site IIIa
Site IIIb
E
L185
T183
D46
P44
W147
R151
R40
N60
E55
F134
S137
M161
L15
E59
W157

**Supplementary Figure 5:** Comparison of the IL-11 signalling complex with the IL-6[4] (PDB ID: 1P9M) and LIF[5] (PDB ID: 1PVH) signalling complexes. A) The site-II binding surface on gp130 for i) IL-11/IL-11Rα, ii) IL-6/IL-6Rα, iii) LIF. B) Interactions between gp130 and the C-helix of i) IL-11, ii) IL-6 and iii) LIF. C) Interactions between gp130 and the N-terminus of i) IL-11, ii) IL-6 and iii) LIF. D) Comparison of the site-III interfaces between i) IL-11/IL-11Rα and ii) IL-6/IL-6Rα. E) Molecular details of the site-III interface of i) IL-11/IL-11R and ii) IL-6/IL-6Rα.

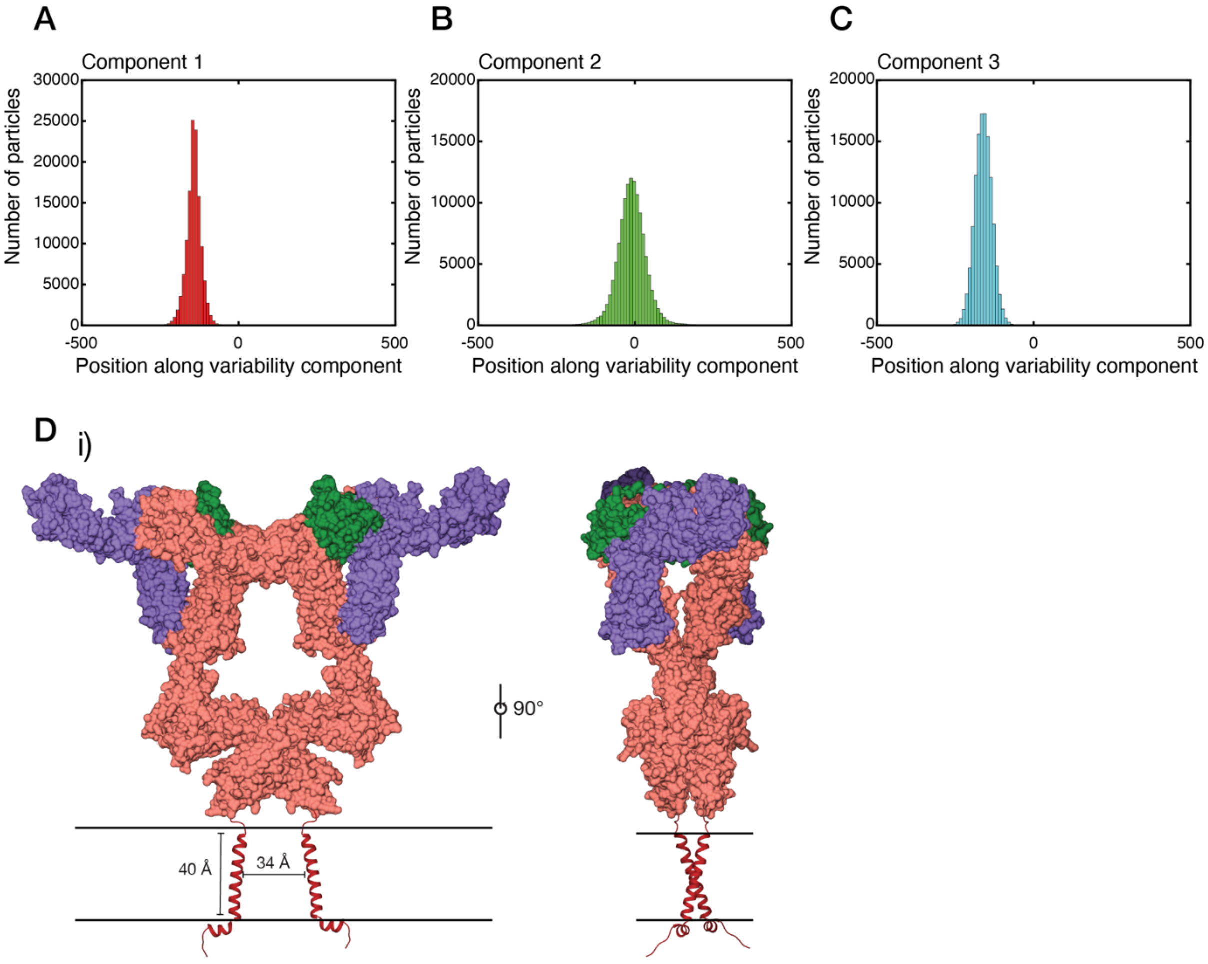

**Supplementary Figure 6:** Histograms indicating the number of particles along each variability component for, A) variability component 1, B) variability component 2, C) variability component 3. D) Model of the membrane-bound IL-11 complex, generated using the Alphafold[6] prediction of full-length gp130 (AF-P40189-F1). The putative position of the cell membrane is indicated using two solid black lines.

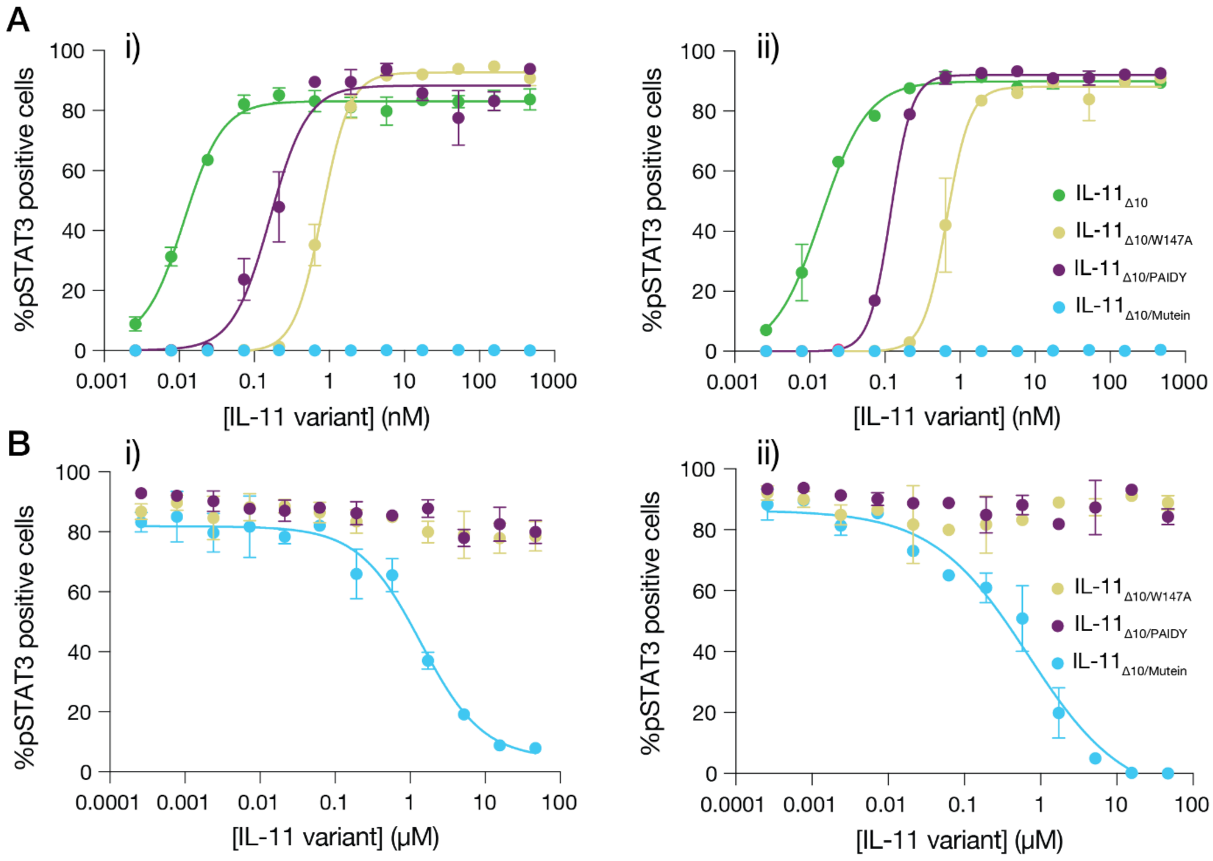

**Supplementary Figure 7:** Replicate dose-response curves for, A) determination of the EC50 of IL-11$_{\Delta 10}$ and other IL-11 variants and B) determination of the IC50 of IL-11$_{\Delta 10/\mathrm{Mutein}}$. Data are presented as standard error of the mean.

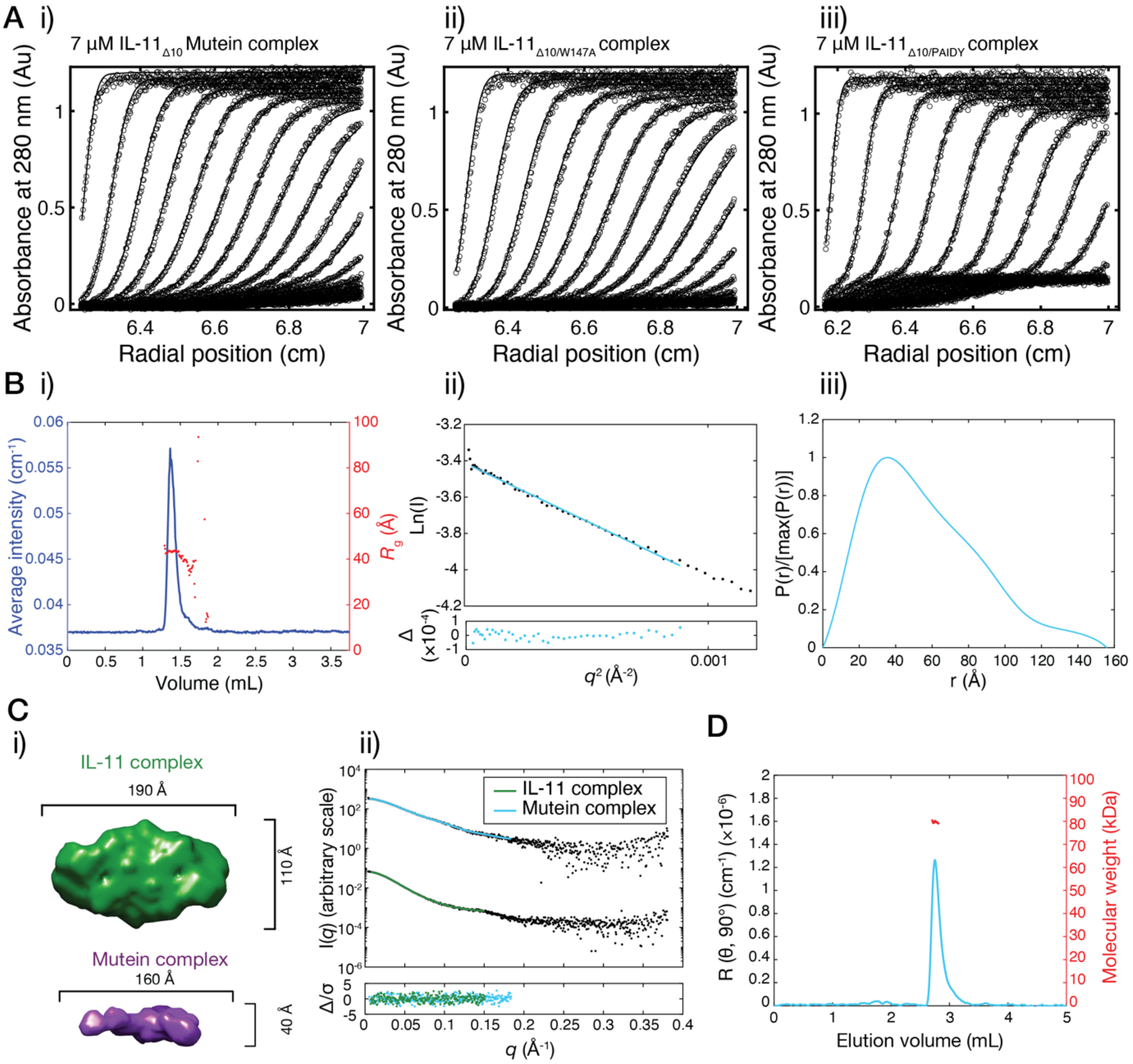

**Supplementary Figure 8:** Raw SV-AUC scans, supplementary SAXS and MALS data for Figure 5. A) Raw SV-AUC scans for the data shown in, i) Figure 5Ai, ii) Figure 5Aii, iii) Figure 5Aiii. B) Supplemental SAXS data for the IL-11$_{\Delta10}$ Mutein/IL-11R$\alpha_{D1\text{-}D3}$/gp130$_{D1\text{-}D3}$ complex, i) SEC-SAXS chromatogram, ii) Guinier plot, iii) pairwise distance distribution (P(r)) plot). C) *Ab inito* SAXS modelling of the complexes, calculated using *DAMMIN*[7], between IL-11$_{\Delta10}$/IL-11$_{\Delta10}$ Mutein, gp130$_{D1\text{-}D3}$ and IL-11R$\alpha_{D1\text{-}D3}$, i) shows the models, and ii) shows the fit to the raw scattering data. D) MALS data for the IL-11$_{\Delta10}$ Mutein/IL-11R$\alpha_{D1\text{-}D3}$/gp130$_{D1\text{-}D3}$ complex.

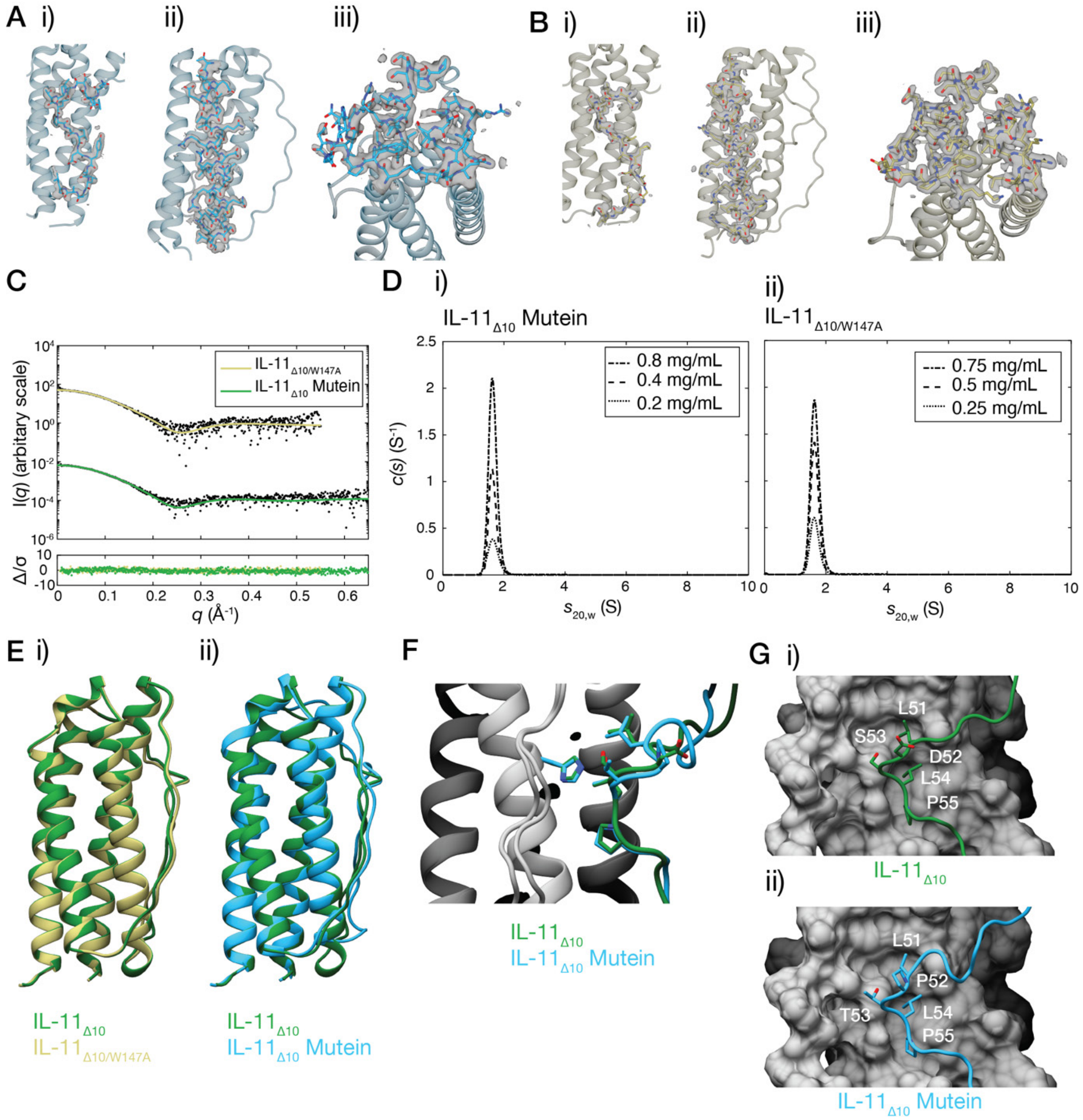

**Supplementary Figure 9:** Representative electron density, additional structural representations and biophysical characterisation of IL-11$_{\Delta10}$ Mutein and IL-11$_{\Delta10/W147A}$. A) Representative electron density for IL-11$_{\Delta10}$ Mutein, i) the AB loop, including the mutated PAIDY sequence, ii) helix C, iii) the site-III interface, including A147. B) Representative electron density for IL-11$_{\Delta10/W147A}$, i) the AB loop, including the AMSAG sequence, ii) helix C, iii) the site-III interface, including A147. C) SAXS data collected on IL-11$_{\Delta10}$ Mutein and IL-11$_{\Delta10/W147A}$, the fit shown is to the crystal structure coordinates. D) Continuous sedimentation coefficient (c(s)) distributions for i) IL-11$_{\Delta10}$ Mutein and ii) IL-11$_{\Delta10/W147A}$, each measured at three concentrations. For raw scans, see Supplementary Figure 10. E) Overlay of the crystal structure of IL-11$_{\Delta10}$[8] (PDB ID: 6O4O) and the structure of i) IL-11$_{\Delta10/W147A}$, ii) IL-11$_{\Delta10}$ Mutein. F) Overlay of the Ser53/Thr56 region in IL-11$_{\Delta10}$ and IL-11$_{\Delta10}$ Mutein. G) Surface representation of the contacts between the AB loop and the α-helical core of i) IL-11$_{\Delta10}$, ii) IL-11$_{\Delta10}$ Mutein.

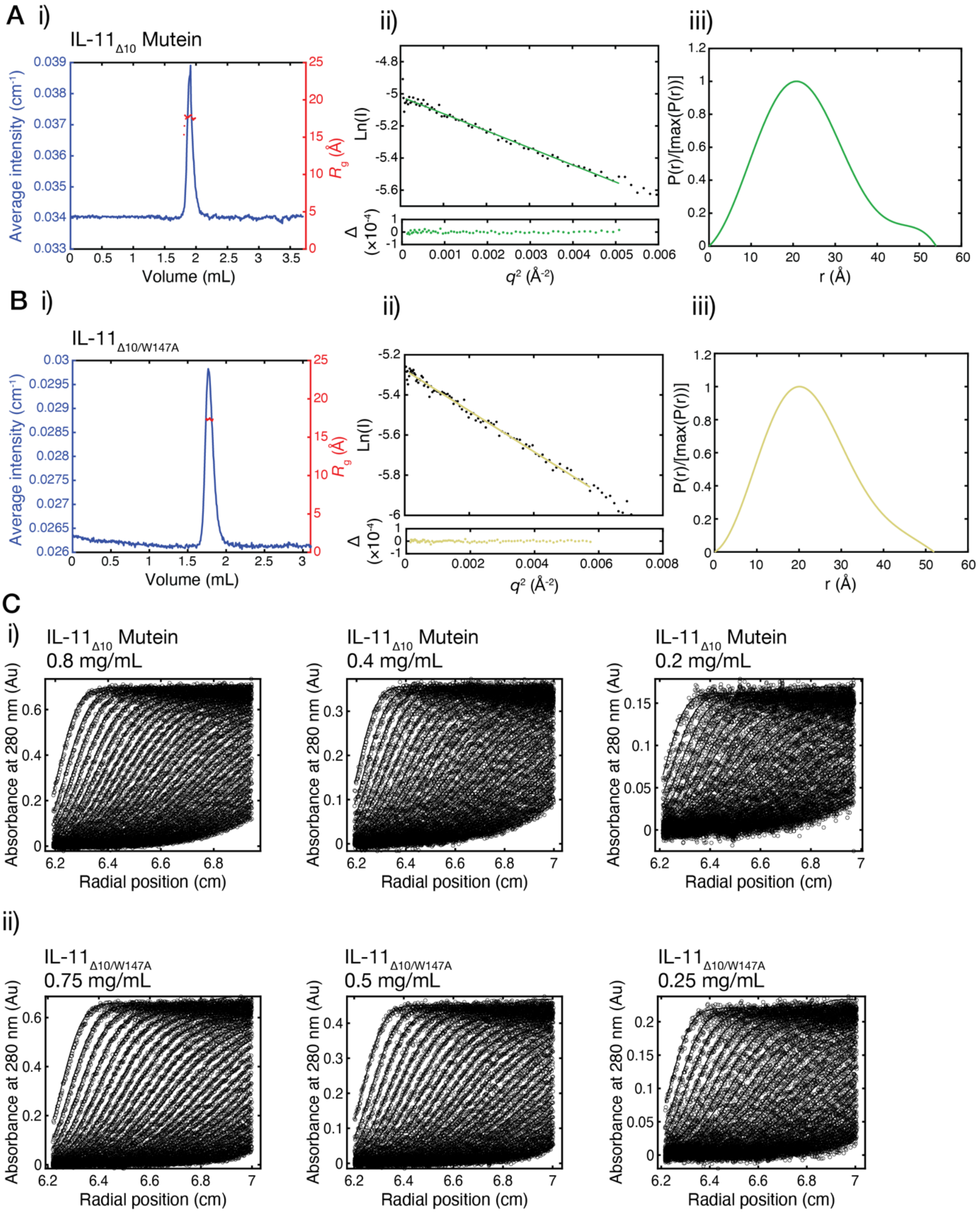

**Supplementary Figure 10:** Supplementary SAXS and AUC data, related to Supplementary Figure 9C-D. A) Supplemental SAXS data for IL-11$_{\Delta10}$ Mutein, i) SEC-SAXS chromatogram, ii) Guinier plot, iii) pairwise distance distribution (P(r)) plot). A) Supplemental SAXS data for IL-11$_{\Delta10/W147A}$, i) SEC-SAXS chromatogram, ii) Guinier plot, iii) pairwise distance distribution (P(r)) plot). C) Raw SV-AUC scans for the data shown in, i) Supplementary Figure 9Di, ii) Supplementary Figure 9Dii.

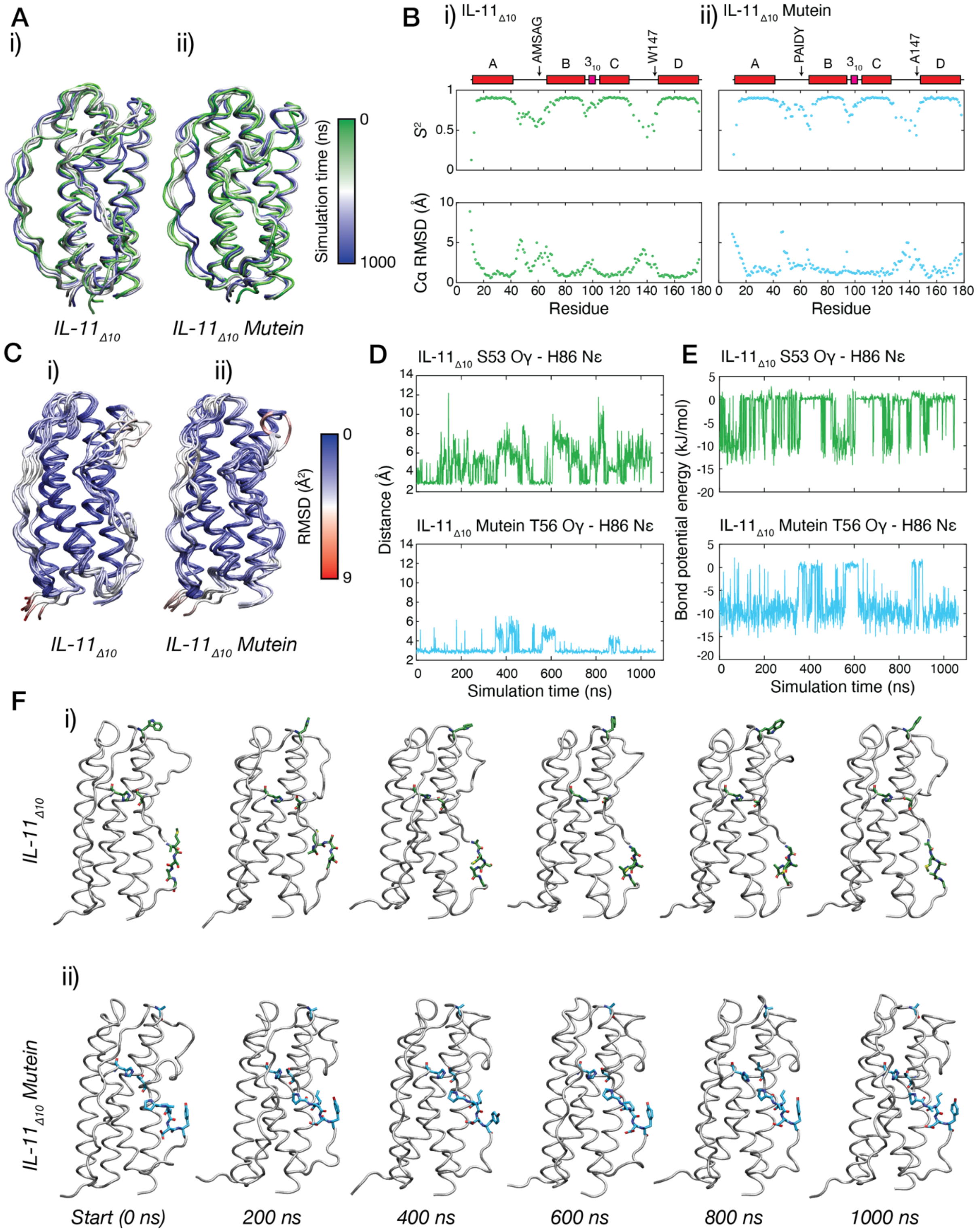

**Supplementary Figure 11:** Supplementary MD data, and DSF data. A) Overlay of frames from the 1 μs MD simulation of i) IL-11$_{\Delta10}$ and ii) IL-11$_{\Delta10}$ Mutein. Frames are shown at 200 ns intervals. B) Order parameter ($S^2$) and Cα RMSD values calculated from the 1 μs MD simulation of i) IL-11$_{\Delta10}$ and ii) IL-11$_{\Delta10}$ Mutein. A schematic representation of the secondary structure is shown above the plot, and the location of the mutations is indicated. C) Overlay of frames from the 1 μs MD simulation of i) IL-11$_{\Delta10}$ and ii) IL-11$_{\Delta10}$ Mutein, coloured by Cα RMSD. Frames are shown at 200 ns intervals. D) Distance between the γ oxygen of T56/S53 in IL-11$_{\Delta10}$ or IL-11$_{\Delta10}$ Mutein through a 1 μs MD simulation. D) Estimated hydrogen bond potential energy for the S/T Oγ and H Nε for IL-$11_{\Delta10}$ or IL-$11_{\Delta10}$ Mutein through a 1 µs MD simulation. F) 200 ns snapshots of the MD simulations of i) IL-$11_{\Delta10}$ and ii) IL-$11_{\Delta10}$ Mutein. The AMSAG/PAIDY sequence, S53/T56, H86 and W147/A147 residues are displayed in the figure.

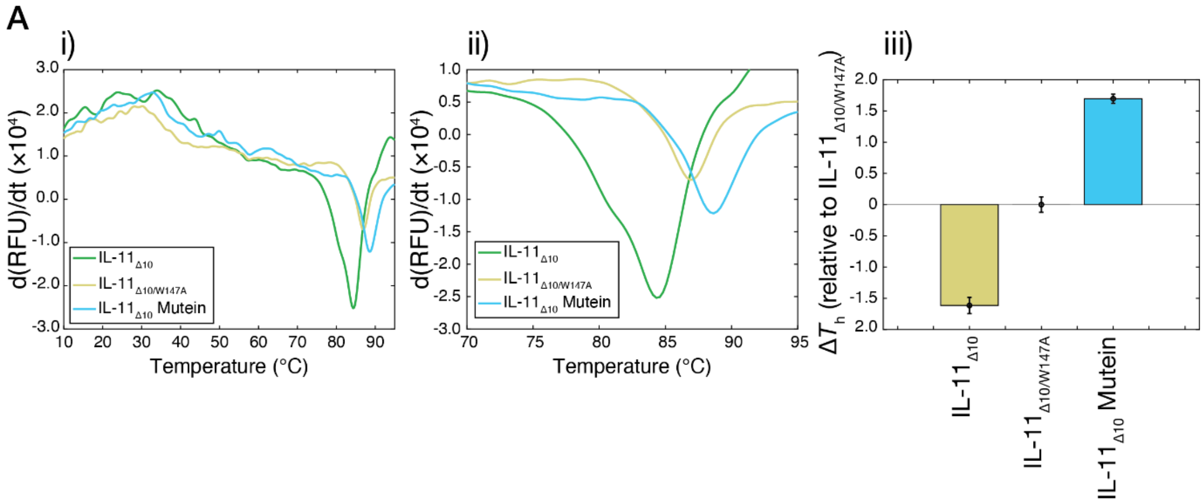

**Supplementary Figure 12:** Differential scanning fluorometry thermal melt data. A) Representative DSF first-derivative melt curve, i) with ii) highlighting the region of interest, iii) Bar graph, showing $\Delta T_h$ (relative to IL-11$_{\Delta 10/W147A}$), error bars indicate standard error, n = 3 independent experiments.

**Supplementary Tables:**

**Supplementary Table 1:** Cryo-EM data collection, data processing and model building statistics.

| | *IL-11 signalling complex gp130$_{D1\text{-}D3}$, IL-11$_{\Delta 10}$, IL-11Rα$_{D1\text{-}D3}$* | *IL-11 signalling complex gp130$_{EC}$, IL-11$_{\Delta 10}$, IL-11Rα$_{D1\text{-}D3}$* |
|---|---|---|
| **Data collection and image processing** | | |
| Magnification | 100,000 | 100,000 |
| Electron energy (kV) | 200 | 200 |
| Electron exposure ($e^-$/Å$^2$) | 52 | 52 |
| Defocus range (μm) | 0.8-2.0 | 0.8-2.0 |
| Pixel size (Å) | 1.31 | 1.31 |
| Starting model | *De novo* | *De novo* |
| Symmetry imposed | C2 | C2 |
| Total number of micrographs | 2010 | 6,861 |
| Initial particle images | 3,082,563 | 694,360 |
| Final particle images | 204,455 | 125,373 |
| Map resolution (Å) | 3.5 | 3.76 |
| FSC threshold | 0.143 | 0.143 |
| **Model building and refinement** | | |
| Initial models used | PDB IDs 6O4O[8], 1I1R[9] (chain A), unpublished structure of IL-11Rα | PDB IDs 6O4O[8], 1I1R[9] (chain A), 3L5I[10], unpublished structure of IL-11Rα |
| Model resolution (Å) | 3.9 | 4.1 |
| FSC threshold | 0.5 | 0.5 |
| Sharpening *B* factor (Å$^2$) | 151.24 | 130.98 |
| *Model composition* | | |
| Non-hydrogen atoms | 10610 | 13550 |
| Amino acid residues | 1326 | 1714 |
| Protein molecules | 10 | 10 |
| *Real-space correlation* | | |
| CCvolume | 0.72 | 0.72 |
| CCmask | 0.73 | 0.74 |
| Mean *B* factor (Å$^2$) | 54.86 | 72.88 |
| RMS deviations | | |
| Bond lengths (Å) (outliers > 4σ) | 0.007 (0) | 0.010 (0) |
| Bond angles (°) (outliers > 4σ) | 1.386 (55) | 1.383 (48) |
| *Validation* | | |
| *MolProbity* score | 2.16 | 2.33 |

| | | |
|---|---|---|
| Clashscore | 18.79 | 23.97 |
| Rotamer outliers (%) | 0.52 | 0.00 |
| CaBLAM outliers (%) | 1.70 | 2.49 |
| Cβ outliers | 0.32 | 0.25 |
| *Ramachandran plot* | | |
| Favoured (%) | 94.19 | 92.83 |
| Allowed (%) | 5.66 | 6.46 |
| Outliers (%) | 0.15 | 0.71 |

**Supplementary Table 2**: X-ray crystallography data collection and refinement statistics.

| | *IL-11 signalling complex $gp130_{D1\text{-}D3}$, $IL\text{-}11_{FL}$, $IL\text{-}11R\alpha_{EC}$* | *$IL11_{\Delta10}$ Mutein* | *$IL11_{\Delta10/W147A}$* |
|---|---|---|---|
| **Data collection** | | | |
| Space group | *P* $3_1$ 1 2 | *P* $2_1$ | *P* $2_1$ $2_1$ 2 |
| Wavelength (Å) | 0.9537 | 0.9537 | 0.9537 |
| Number of images | 3600 | 3600 | 3600 |
| Oscillation range per image (˚) | 0.1 | 0.1 | 0.1 |
| Detector | Eiger 16M | Eiger 16M | Eiger 16M |
| Cell dimensions | | | |
| *a*, *b*, *c* (Å) | 163.41, 163.41, 506.62 | 27.23, 37.097, 68.532 | 38.505, 133.996, 27.087 |
| α, β, γ (˚) | 90, 90, 120 | 90, 101.306, 90 | 90, 90, 90 |
| Resolution (Å) | 49.275-3.779 (4.197-3.779) | 37.10-1.80 (1.841.80) | 44.67-1.48 (1.51-1.48) |
| $R_{sym}$[†] | 0.286 (2.522) | 0.07 (1.167) | 0.082 (1.859) |
| $R_{meas}$[§] | 0.294 (2.584) | 0.083 (1.374) | 0.089 (2.003) |
| $R_{pim}$[‡] | 0.064 (0.558) | 0.043 (0.719) | 0.034 (0.743) |
| $CC_{1/2}$ | 0.998 (0.548) | 0.999 (0.603) | 0.999 (0.590) |
| $I/\sigma(I)$ | 10.6 (1.5) | 11.8 (1.4) | 12.9 (1.3) |
| Total observations | 947761 (48027) | 85885 (5234) | 313134 (16025) |
| Unique reflections | 45270 (2262) | 12637 (761) | 24354 (1182) |
| Completeness (%) | | 100.0 (100.0) | 100.0 (100.0) |
| Spherical (%) | 58.1 (10.8) | | |
| Ellipsoidal (%) | 92.8 (72.0) | | |
| Multiplicity | 20.9 (21.2) | 6.8 (6.9) | 12.9 (13.6) |
| Wilson *B* factor (Å$^2$) | 126.31 | 25.604 | 23.185 |
| **Refinement** | | | |
| Resolution (Å) | 40.86-3.78 (3.91-3.78) | 33.61-1.80 (1.86-1.80) | 37.02-1.48 (1.53-1.48) |

| | | | |
|---|---|---|---|
| Reflections used in refinement | 43181 (296) | 12627 (1241) | 23081 (2232) |
| $R_{free}$ reflections | 2054 (21) | 1173 (68) | 1216 (129) |
| $R_{work}$ | 0.2805 (0.4323) | 0.1978 (0.2899) | 0.1889 (0.2732) |
| $R_{free}$ | 0.2966 (0.4378) | 0.2306 (0.2773) | 0.2185 (0.2696) |
| Protein molecules in asymmetric unit | 18 | 1 | 1 |
| Total nonhydrogen atoms | 35652 | 1390 | 1429 |
| Protein | 35652 | 1304 | 1308 |
| Ligand/ion | 0 | 5 | 6 |
| Solvent | 0 | 71 | 114 |
| *Mean* B *factor (Å$^2$)* | 176.20 | 46.65 | 37.48 |
| Protein | 176.20 | 46.81 | 37.10 |
| Ligand/ion | N/A | 59.96 | 49.03 |
| *RMS deviations* | | | |
| Bond lengths (Å) (outliers > 4σ) | 0.005 (6) | 0.003 (0) | 0.005 (0) |
| Bond angles (°) (outliers > 4σ) | 1.156 (86) | 0.630 (0) | 0.791 (0) |
| Rotamer outliers | 0.16 | 0.73 | 0.00 |
| Clashscore | 14.16 | 4.14 | 4.10 |
| Cβ outliers | 0 | 0 | 0 |
| *Molprobity* score | 2.06 | 1.20 | 1.19 |
| *Ramachandran Plot* | | | |
| Favoured (%) | 94.10 | 98.20 | 98.80 |
| Allowed (%) | 5.22 | 1.80 | 1.20 |
| Outliers (%) | 0.68 | 0.0 | 0.0 |

† $R_{sym} = \sum_{hkl}\sum_i |I_i(hkl) \quad \langle I(hkl)\rangle| / \sum_{hkl}\sum_i I_i(hkl)$

§ $R_{meas} = \sum_{hkl} [N/(N \quad 1)]^{1/2} \sum_i |I_i(hkl) \quad \langle I(hkl)\rangle| / \sum_{hkl}\sum_i I_i(hkl)$

‡ $R_{\mathrm{pim}} = \sum_{hkl} [1/(N \quad 1)]^{½} \sum_{i} |I_{i} (hkl) \quad \langle I (hkl) \rangle \ | \ / \sum_{hkl} \sum_{i} I_{i} (hkl)$

$CC_{1/2}$ = Pearson correlation coefficient between independently merged half datasets

**Supplementary Table 3**: SAXS data collection and refinement statistics.

| | $gp130_{D1\text{-}D3}$ complex [a] | $gp130_{EC}$ complex [b] | $gp130_{D2\text{-}D3}$ complex [c] | $IL11_{\Delta 10}$ Mutein complex [d] | $IL11_{\Delta 10}$ Mutein | $IL11_{\Delta 10/W147A}$ |
|---|---|---|---|---|---|---|
| **SAXS data collection** | | | | | | |
| Instrument/source | Australian Synchrotron SAXS/WAXS beamline equipped with Pilatus 2M detector and sheathflow cell for SEC-SAXS[11,12]. | | | | | |
| Wavelength (Å) | 1.078 | | | | | |
| Beam energy (keV) | 11.5 | | | | | |
| Beam size (µm) | 250 × 130 | | | | | |
| Sample-to-detector distance (mm) | 3538 | 2210 | 3538 | 3538 | 2038 | 2210 |
| *q* measurement range (Å$^{1}$) [a] | 0.004-0.38 | 0.005-0.55 | 0.004-0.38 | 0.004-0.38 | 0.007-0.664 | 0.005-0.55 |
| Absolute scaling method | Comparison with scattering from 1 mm pure water | | | | | |
| Normalization | To transmitted intensity from beamstop counter | | | | | |
| Exposure time | 1 s measurements from SEC-SAXS elution | | | | | |
| Sample temperature (K) | 293 | | | | | |
| **SEC-SAXS parameters** | | | | | | |
| Column | Superdex 200 5/150 Increase | | | | | |
| Flow rate (mL/min) | 0.45 | | | | | 0.4 |
| Loading concentration (mg/mL) | 2 | | | | 5 | |
| Injection volume (µL) | 50 | | | | | |
| Solvent | 20 mM Tris-HCl pH 8.5, 150 mM NaCl, 0.2% sodium azide | | | | | |
| **Software employed** | | | | | | |
| SAXS data reduction | *I*(*q*) vs *q* using Scatterbrain 2.8.2, SECSAXS solvent subtraction using *CHROMIXS* from *ATSAS* 2.8.3 | | | | | |

| | | | | | | |
|---|---|---|---|---|---|---|
| Basic analysis (Guinier, *P(r)*, molecular mass) | *PRIMUS* from *ATSAS* 2.8.3, *GNOM* from *ATSAS* 2.83 | | | | | |
| Shape modelling | *DAMMIF* from *ATSAS* 2.8.3, *DAMAVER* from *ATSAS* 2.8.3, *DAMMIN* from *ATSAS* 2.8.3 | | | | | |
| Calculation of theoretical intensities | *CRYSOL* from *ATSAS* 2.8.3 | | | | | |
| **Structural parameters** | | | | | | |
| Mass from $V_c$ (kDa) (expected mass, ratio to expected, in brackets) [b] | 182.2 (169.8, 0.93) | 290.0 (234.8, 0.81) | 81.6 (73.6, 0.90) | 87.9 (84.9, 0.96) | 16.7 (18.2, 0.91) | 16.7 (18.2, 0.91) |
| **Guinier analysis** | | | | | | |
| $R_g$ (Å) | 53.10 ± 0.19 | 61.99 ± 0.46 | 36.49 ± 0.14 | 43.41 ± 0.18 | 17.82 ± 0.13 | 17.36 ± 0.12 |
| *I(0)* ($cm^{1}$) | $0.068 \pm 1.4\times10^{-4}$ | $0.039 \pm 2.2\times10^{-4}$ | $0.029 \pm 7\times10^{-5}$ | $0.033 \pm 8.4\times10^{-5}$ | $0.0066 \pm 2.6\times10^{-5}$ | $0.0051 \pm 2.1\times10^{-5}$ |
| $qR_g$ min,max | 0.41, 1.22 | 0.42, 1.32 | 0.24, 1.31 | 0.23, 1.29 | 0.20, 1.27 | 0.2, 1.32 |
| *P(r)* analysis [c] | | | | | | |
| $R_g$ (Å) | 54.08 ± 0.13 | 62.66 ± 0.37 | 37.52 ± 0.15 | 45.24 ± 0.19 | 17.80 ± 0.88 | 17.33 ± 0.69 |
| *I(0)* ($cm^{1}$) | $0.0677 \pm 1.2\times10^{-4}$ | $0.03891 \pm 1.9\times10^{-4}$ | $0.02908 \pm 6.8\times 10^{-5}$ | $0.033 \pm 9.7\times10^{-5}$ | $0.0066 \pm 2.2 \times 10^{-5}$ | $0.0051 \pm 1.6 \times 10^{-5}$ |
| $D_{max}$ (Å) | 176 | 210 | 133 | 156 | 54 | 52 |
| Porod volume ($Å^3$) | 404000 | 709000 | 127000 | 159000 | 20000 | 22200 |
| **Shape modelling** | | | | | | |
| *DAMMIF* (10 calculations, default parameters) | | | | | | |
| *q* range for fitting (Å) | 0.007-0.15 | | | 0.00054 – 0.18 | | |

| | | | | | | |
|---|---|---|---|---|---|---|
| Symmetry, anisotropy assumptions | *P*2, none | | | *P*1, none | | |
| Constant adjustment to intensities | $5.11 \times 10^{-4}$ | | | $1.67 \times 10^{-4}$ | | |
| NSD (standard deviations) | 1.281 (0.176) | | | 1.028 (0.081) | | |
| $\chi^2$ range | 1.114-1.129 | | | 0.9981.018 | | |
| Resolution (from *SASRES*[13]) (Å) | 50 ± 4 | | | 47 ± 4 | | |
| *DAMMIN* (default parameters) | | | | | | |
| *q* range for fitting (Å) | 0.007-0.15 | | | 0.00054 – 0.18 | | |
| Symmetry, anisotropy assumptions | *P2*, none | | | *P*1, none | | |
| $\chi^2$ | 1.009 | | | 0.975 | | |
| Constant adjustment to intensities | $5.03 \times 10^{-4}$ | | | $1.58 \times 10^{-4}$ | | |
| **Atomic modelling** | | | | | | |
| *CRYSOL* (no constant subtraction) | | | | | | |
| Structure | gp130$_{EC}$ complex residues 2-300 chain A, D, chain B, E, chain C, F | gp130$_{EC}$ complex with gp130 D5-D6 domains modelled | gp130$_{EC}$ complex residues 100-300 chain A, chain B, chain C | gp130$_{EC}$ complex residues 2-269 chain A, chain B, chain C | IL-11$_{\Delta10/\mathrm{Mutein}}$ crystal structure | IL-11$_{\Delta10/\mathrm{W147A}}$ crystal structure |
| $\chi^2$ | 3.06 | 2.66 | 1.76 | 1.72 | 1.37 | 0.91 |
| Calculated $R_g$ (Å) | 53.38 | 61.04 | 36.12 | 43.30 | 17.61 | 17.39 |
| Structure | IL-11$_{\Delta10}$ complex crystal structure chains A-F | | | | | |
| $\chi^2$ | 2.32 | | | | | |

| | |
|---|---|
| Calculated $R_g$ (Å) | 54.00 |

[a] Hexameric complex between gp130$_{D1\text{-}D3}$/IL-11$_{\Delta 10}$/IL-11Rα$_{D1\text{-}D3}$

[b] Hexameric complex between gp130$_{EC}$/IL-11$_{\Delta 10}$/IL-11Rα$_{D1\text{-}D3}$

[c] Trimeric complex between gp130$_{D2\text{-}D3}$/IL-11$_{\Delta 10}$/IL-11Rα$_{D1\text{-}D3}$

[d] Trimeric complex between gp130$_{D1\text{-}D3}$/IL-11$_{\Delta 10/Mutein}$/IL-11Rα$_{D1\text{-}D3}$

**Supplementary Table 4**: Complete ITC thermodynamic data. Errors are ± standard error, n = 3 independent titrations for all experiments, errors shown are ± standard error.

| | | $K_D$ (nM) | ΔH (kJ/mol) | ΔS (J/molK) | ΔG (kJ/mol) | Incompetent receptor fraction [a] | T (K) |
|---|---|---|---|---|---|---|---|
| IL-$11_{\Delta10}$/IL-$11R\alpha_{D1\text{-}D3}$ | $gp130_{D1\text{-}D3}$ | 3 ± 2 | -34 ± 0.4 | 51 ± 9.4 | -49 ± 3.0 | 0.32 ± 0.008 | 288 |
| IL-$11_{\Delta10}$/IL-$11R\alpha_{D1\text{-}D3}$ | $gp130_{D2\text{-}D3}$ | 380 ± 190 | 27 ± 2.0 | 220 ± 4.2 | -36 ± 1.7 | 0.25 ± 0.03 | 288 |
| IL-$11_{\Delta10}$/IL-$11R\alpha_{D1\text{-}D3}$ | $gp130_{EC}$ | 4 ± 2 | -21 ± 0.7 | 94 ± 5.4 | -49 ± 1.2 | 0.36 ±0.03 | 288 |
| IL-$11_{\Delta10/W147A}$ | IL-$11R\alpha_{D1\text{-}D3}$ | 10 ± 8 | -22 ± 0.6 | 87 ± 11 | -48 ± 2.7 | 0.08 ± 0.04 | 303 |
| IL-$11_{\Delta10}$ Mutein | IL-$11R\alpha_{D1\text{-}D3}$ | 38 ± 9.4 | -25 ± 1.1 | 64 ± 4.1 | -44 ± 0.6 | 0.04 ± 0.02 | 303 |
| IL-$11_{\Delta10/PAIDY}$ | IL-$11R\alpha_{D1\text{-}D3}$ | 81 ± 44 | -30 ± 2.2 | 38 ± 11 | -42 ± 1.4 | 0.15 ± 0.05 | 303 |
| IL-$11_{\Delta10}$ Mutein/IL-$11R\alpha_{D1\text{-}D3}$ | $gp130_{D1\text{-}D3}$ | 55 ± 4.1 | 22 ± 0.7 | 220 ± 1.9 | -40 ± 0.2 | 0.43 ± 0.03 | 288 |
| IL-$11_{\Delta10/W147A}$/IL-$11R\alpha_{D1\text{-}D3}$ | $gp130_{D2\text{-}D3}$ | 130 ± 14 | 24 ± 0.7 | 210 ± 2.6 | -38 ± 0.2 | 0.33 ± 0.03 | 288 |
| IL-$11_{\Delta10/PAIDY}$/IL-$11R\alpha_{D1\text{-}D3}$ | $gp130_{D2\text{-}D3}$ | 60 ± 16 | 23 ± 0.8 | 220 ± 2.5 | -40 ± 0.7 | 0.35 ± 0.02 | 288 |
| IL-$11_{\Delta10/PAIDY}$/IL-$11R\alpha_{D1\text{-}D3}$ | $gp130_{D1\text{-}D3}$ | 20 ± 17 | -31 ± 3.0 | 51 ± 6.4 | -47 ± 2.3 | 0.34 ± 0.07 | 288 |

[a] Similar to (1-N), see ref [14].

**Supplementary Table 5:** Complete SPR thermodynamic and kinetic data, n = 2 experiments, errors shown are ± standard error.

| | | $K_D$ (nM) | $k_d$ ($\times 10^{-1}$ $s^{-1}$) | $k_a$ ($\times 10^{6}$ $M^{-1}$ $s^{-1}$) | T (K) |
|---|---|---|---|---|---|
| $IL11_{\Delta 10}$ - Avitag | IL-11R$\alpha_{D1\text{-}D3}$ | 78 ± 1 | 1.1 ± 0.2 | 1.4 ± 0.2 | 298 |
| $IL11_{\Delta 10}$ Mutein - Avitag | IL-11R$\alpha_{D1\text{-}D3}$ | 33 ± 3 | 0.34 ± 0.01 | 1.0 ± 0.1 | 298 |

**Supplementary Discussion**

Gp130 is a receptor shared by most other members of the IL-6 family of cytokines. Structures have been solved of the hexameric IL-6 signalling complex[4], which forms analogous site-II and site-III interactions with gp130, and the LIF/gp130 complex, which forms an analogous site-II interaction with gp130[5] (Supplementary Figure 5). The LIF and IL-6 gp130 site-II interactions have been previously compared[5]. IL-11, IL-6 and LIF interact with a similar surface on gp130 (Supplementary Figure 5A), with major common contacts are formed by residues 142-147 and 165-171 on gp130, the N-terminal region of the cytokine and B helix of the cytokine. F163 of gp130 forms the major contact to the N-terminal end of the helix of the cytokine. The cytokine B-helix contacts a similar region of gp130 in the three structures, however in IL-11 the contacts are more extensive, a consequence of the additional surface area buried by arginine residues 111, 114, 117 and 118 in IL-11 (Supplementary Figure 5Bi). The contacts with the B-helix of IL-6 and LIF are dominated by a number of small hydrophobic residues (Supplementary Figure 5Bii-iii), and are overall less extensive compared to IL-11. The N-terminal region of IL-11, IL-6 and LIF also interact with gp130 (Supplementary Figure 5C). The N-terminus of LIF forms more extensive contacts with gp130 compared to IL-6 and IL-11, which may underpin the ability of LIF to interact with gp130 without first interacting with an α-receptor[5] (Supplementary Figure 5Cii). The interaction between IL-6 and IL-11 with gp130 is complimented by an additional site-IIB interface between the α-receptor and gp130. In both complexes, the surface bound on gp130 is very similar (Supplementary Figure 5A), with the binding surface predominantly formed by residues 250-265 of gp130. Notably, the interaction between IL-11Rα and gp130 is more electrostatic in character and results in formation of ten hydrogen bonds, compared to five for the IL-6Rα/gp130 complex.

IL-6 and IL-11 both form an additional interaction with D1 of gp130 at site-III to form the hexameric signalling complex. LIF forms an analogous interaction with D4 of LIFR[15]. The IL-6/gp130 site-III interaction is more extensive compared to the IL-11/gp130 site-III interaction (Supplementary Figure 5D-E). The surface bound on D1 of gp130 is similar between the two cytokines (Supplementary Figure 5D). In both interactions, the predominant contacts are made by the N-terminal end of the AB loop and the D helix of the cytokine, the N-terminus of gp130, and residues 90-98 of gp130. IL-6 makes more extensive contacts between the N-terminal end of the AB loop and the N-terminus of gp130 (Supplementary Figure 5Eii). Analogous interactions are not present in the IL-11 complex; indeed, the N-terminal loop of gp130 is poorly resolved in all of our density maps. An analogous tryptophan (W147 in IL-11, W157 in IL-6), at the N-terminal end of the D helix forms a key site-III contact in both complexes (Supplementary Figure 5E). Additional hydrophobic residues present in the IL-6 complex at the end of the D helix (e.g. L158) that are not present in the IL-11 complex. Similarly, the site-IIIb interface is significantly less extensive in the IL-11 complex, compared to the IL-6 complex (Supplementary Figure 5E). In both complexes the major contact between gp130 and the α-receptor is formed by a short loop in gp130 (residues 86-89), however in the IL-6 complex these contacts are more extensive, and additional contacts are contributed by an additional short loop (residues 35-37). The altered pose of IL-11Rα compared to IL-6Rα in the complex appears to have reduced the contacts formed between IL-11Rα and gp130 at site-IIIb (Supplementary Figure 5E). Overall, the IL-11 complex structure reinforces a previous suggestion[5] that gp130 has evolved to interact with unique and structurally diverse cytokines, in both the CHR and D1 of gp130.

**References:**

1 Adams, P. D. *et al.* PHENIX: A comprehensive Python-based system for macromolecular structure solution. *Acta Crystallographica Section D: Biological Crystallography* **66**, 213-221, doi:10.1107/S0907444909052925 (2010).

2 Afonine, P. V. *et al.* New tools for the analysis and validation of Cryo-EM maps and atomic models. *Acta Crystallographica Section D Structural Biology* **74**, 814-840, doi:10.1101/279844 (2018).

3 Kucukelbir, A., Sigworth, F. J. & Tagare, H. D. Quantifying the local resolution of cryo-EM density maps. *Nature Methods* **11**, 63-65, doi:10.1038/nmeth.2727 (2014).

4 Boulanger, M. J., Chow, D.-c., Brevnova, E. E. & Garcia, K. C. Hexameric structure and assembly of the interleukin-6/IL-6 alpha-receptor/gp130 complex. *Science* **300**, 2101-2104, doi:10.1126/science.1083901 (2003).

5 Boulanger, M. J., Bankovich, A. J., Kortemme, T., Baker, D. & Garcia, K. C. Convergent mechanisms for recognition of divergent cytokines by the shared signaling receptor gp130. *Molecular Cell* **12**, 577-589, doi:10.1016/S1097-2765(03)00365-4 (2003).

6 Jumper, J. *et al.* Highly accurate protein structure prediction with AlphaFold. *Nature* **596**, 583-589, doi:10.1038/s41586-021-03819-2 (2021).

7 Svergun, D. Restoring low resolution structure of biological macromolecules from solution scattering using simulated annealing. *Biophysical Journal* **76**, 2879-2886 (1999).

8 Metcalfe, R. D. *et al.* The structure of the extracellular domains of human interleukin 11 α-receptor reveals mechanisms of cytokine engagement. *Journal of Biological Chemistry (DOI: 10.1074/jbc.RA119.012351)*, doi:10.1074/jbc.RA119.012351 (2020).

9 Chow, D.-C., He, X., Snow, a. L., Rose-John, S. & Garcia, K. Structure of an extracellular gp130 cytokine receptor signaling complex. *Science* **291**, 2150-2155, doi:10.1126/science.1058308 (2001).

10 Xu, Y. *et al.* Crystal structure of the entire ectodomain of gp130: Insights into the molecular assembly of the tall cytokine receptor complexes. *Journal of Biological Chemistry* **285**, 21214-21218, doi:10.1074/jbc.C110.129502 (2010).

11 Ryan, T. M. *et al.* An optimized SEC-SAXS system enabling high X-ray dose for rapid SAXS assessment with correlated UV measurements for biomolecular structure analysis:. *Journal of Applied Crystallography* **51**, 97-111, doi:10.1107/S1600576717017101 (2018).

12 Kirby, N. *et al.* Improved radiation dose efficiency in solution SAXS using a sheath flow sample environment. *Acta Crystallographica Section D Structural Biology* **72**, 1254-1266, doi:10.1107/S2059798316017174 (2016).

13 Tuukkanen, A. T., Kleywegt, G. J. & Svergun, D. I. Resolution of ab initio shapes determined from small-angle scattering *IUCrJ* **3**, 440-447, doi:10.1107/s2052252516016018 (2016).

14 Zhao, H. & Schuck, P. Combining biophysical methods for the analysis of protein complex stoichiometry and affinity in SEDPHAT. *Acta Crystallographica Section D: Biological Crystallography* **71**, 3-14, doi:10.1107/S1399004714010372 (2015).

15 Huyton, T. *et al.* An unusual cytokine:Ig-domain interaction revealed in the crystal structure of leukemia inhibitory factor (LIF) in complex with the LIF receptor. *Proceedings of the National Academy of Sciences* **104**, 12737-12742, doi:10.1073/pnas.0705577104 (2007).

16 Nandurkar, H. H. *et al.* The human IL-11 receptor requires gp130 for signalling: demonstration by molecular cloning of the receptor. *Oncogene* **12**, 585-593 (1996).

17 Van Den Berg, S., Löfdahl, P. Å., Härd, T. & Berglund, H. Improved solubility of TEV protease by directed evolution. *Journal of Biotechnology* **121**, 291-298, doi:10.1016/j.jbiotec.2005.08.006 (2006).

18 Cabrita, L. D. *et al.* Enhancing the stability and solubility of TEV protease using in silico design. *Protein Science* **16**, 2360-2367, doi:10.1110/ps.072822507 (2007).

19 Kapust, R. B. *et al.* Tobacco etch virus protease: mechanism of autolysis and rational design of stable mutants with wild-type catalytic proficiency. *Protein Engineering* **14**, 993-1000, doi:10.1093/protein/14.12.993 (2001).

20 Putoczki, T. L., Dobson, R. C. J. & Griffin, M. D. W. The structure of human interleukin-11 reveals receptor-binding site features and structural differences from interleukin-6. *Acta Crystallographica Section D-Biological Crystallography* **3**, 2277-2285, doi:10.1107/S1399004714012267 (2014).

21 Kimanius, D., Forsberg, B. O., Scheres, S. H. & Lindahl, E. Accelerated cryo-EM structure determination with parallelisation using GPUs in RELION-2. *Elife* **5**, doi:10.7554/eLife.18722 (2016).

22 Scheres, S. H. Semi-automated selection of cryo-EM particles in RELION-1.3. *J Struct Biol* **189**, 114-122, doi:10.1016/j.jsb.2014.11.010 (2015).

23 Punjani, A., Rubinstein, J. L., Fleet, D. J. & Brubaker, M. A. CryoSPARC: Algorithms for rapid unsupervised cryo-EM structure determination. *Nature Methods* **14**, 290-296, doi:10.1038/nmeth.4169 (2017).

24 Li, X. *et al.* Electron counting and beam-induced motion correction enable near-atomic-resolution single-particle cryo-EM. *Nature Methods* **10**, 584-590, doi:10.1038/nmeth.2472 (2013).

25 Terwilliger, T. C., Sobolev, O. V., Afonine, P. V. & Adams, P. D. Automated map sharpening by maximization of detail and connectivity. *Acta Crystallographica Section D Structural Biology* **74**, 545-559, doi:10.1107/S2059798318004655 (2018).

26 Kabsch, W. Xds. *Acta Crystallographica Section D: Biological Crystallography* **66**, 125-132, doi:10.1107/S0907444909047337 (2010).

27 Evans, P. Scaling and assessment of data quality. *Acta Crystallographica Section D: Biological Crystallography* **62**, 72-82, doi:10.1107/S0907444905036693 (2006).

28 Evans, P. R. & Murshudov, G. N. How good are my data and what is the resolution? *Acta Crystallographica Section D: Biological Crystallography* **69**, 1204-1214, doi:10.1107/S0907444913000061 (2013).

29 McCoy, A. J. *et al.* Phaser crystallographic software *Journal of Applied Crystallography* **40**, 658-674, doi:10.1107/s0021889807021206 (2007).

30 Emsley, P., Lohkamp, B., Scott, W. G. & Cowtan, K. Features and development of Coot. *Acta Crystallographica Section D: Biological Crystallography* **66**, 486-501, doi:10.1107/S0907444910007493 (2010).

31 Afonine, P. V. *et al.* Towards automated crystallographic structure refinement with phenix.refine. *Acta Crystallographica Section D: Biological Crystallography* **68**, 352-367, doi:10.1107/S0907444912001308 (2012).

32 Afonine, P. V. *et al.* Real-space refinement in Phenix for cryo-EM and crystallography. *Acta Crystallographica Section D Structural Biology* **74**, 531-544, doi:10.1101/249607 (2018).

33 Schuck, P. Size-distribution analysis of macromolecules by sedimentation velocity ultracentrifugation and lamm equation modeling. *Biophysical Journal* **78**, 1606-1619, doi:10.1016/S0006-3495(00)76713-0 (2000).
34 Ortega, A., Amorós, D. & García De La Torre, J. Prediction of hydrodynamic and other solution properties of rigid proteins from atomic- and residue-level models. *Biophysical Journal* **101**, 892-898, doi:10.1016/j.bpj.2011.06.046 (2011).
35 Laue, T. M., Shah, B. D., Ridgeway, T. M. & Pelletier, S. L. *Analytical Ultracentrifugation in Biochemistry and Polymer Science*. 90-125 (Cambridge: Royal Society of Chemistry, 1992).
36 Panjkovich, A. & Svergun, D. I. CHROMIXS: automatic and interactive analysis of chromatography-coupled small-angle X-ray scattering data. *Bioinformatics* **34**, 1944-1946
doi:10.1093/bioinformatics/btx846 (2017).
37 Barberato, C. & Koch, M. H. J. CRYSOL - a Program to Evaluate X-ray Solution Scattering of Biological Macromolecules from Atomic Coordinates. *Journal of Applied Crystallography* **28**, 768-773, doi:10.1107/S0021889895007047 (1995).
38 Franke, D. & Svergun, D. I. DAMMIF, a program for rapid ab-initio shape determination in small-angle scattering. *Journal of Applied Crystallography* **42**, 342-346, doi:10.1107/S0021889809000338 (2009).
39 Volkov, V. V. & Svergun, D. I. Uniqueness of ab initio shape determination in small-angle scattering. *Journal of Applied Crystallography* **36**, 860-864, doi:10.1107/S0021889803000268 (2003).
40 Phillips, J. C. *et al.* Scalable molecular dynamics with NAMD. *Journal of Computational Chemistry* **26**, 1781-1802, doi:10.1002/jcc.20289 (2005).
41 Brooks, B. R. *et al.* CHARMM: The Biomolecular Simulation Program. *Journal of Computational Chemistry* **20**, 1545-1614, doi:10.1002/jcc (2009).
42 Pettersen, E. F. *et al.* UCSF Chimera--a visualization system for exploratory research and analysis. *J Comput Chem* **25**, 1605-1612, doi:10.1002/jcc.20084 (2004).
43 Humphrey, W., Dalke, A. & Schulten, K. VMD: Visual Molecular Dynamics. *Journal of Molecular Graphics* **7855**, 33-38, doi:10.1016/0263-7855(96)00018-5 (1996).
